# The human claustrum initiates networks for externally and internally driven task demands

**DOI:** 10.1101/2025.06.26.661853

**Authors:** Brent W. Stewart, Matthew A. Cormie, Michael L. Keaser, Massieh Moayedi, Brian N. Mathur, David A. Seminowicz

## Abstract

Cognitive control is believed to arise from interactions among multiple brain networks depending on task demands. Although several debilitating neuropsychiatric disorders are characterized by cognitive network dysfunction, the neural circuit mechanisms supporting task-dependent network activation are largely unknown. Because the claustrum possesses widespread connections with cortex and can synchronize distant cortical regions, we tested whether the claustrum activates task-dependent network states using fMRI during working memory (*n* = 420) and autobiographical memory (*n* = 35), tasks which elicit opposing responses from key cognitive control networks. In both tasks, the claustrum exhibited increased activity and excitatory influence on task-associated cognitive control network nodes, with corroborating underlying structural connectivity. The claustrum also displayed stronger excitatory effective connectivity during task performance and greater structural connectivity with task-related network nodes than regions prominently implicated in directing network states—the anterior insula and pulvinar. These findings establish a role for the claustrum in initiating network states for cognitive control.

**Significance:** Cognitive functioning is supported by large-scale networks across the brain. Yet, the neural circuit mechanisms supporting task-dependent network activation are largely unknown. Circuit analyses using human functional and structural neuroimaging in this study found that the claustrum, a subcortical nucleus, activates during multiple cognitive tasks eliciting a wide range of network states, possesses strong anatomical connections with cognitive control network nodes, and exerts excitatory influence on task-associated network regions. These results establish the claustrum as a network activator subserving cognitive control.

## Introduction

Cognitive control enables the selection of appropriate actions by filtering large sets of sensory inputs and possible motor responses based on internal goals (Miller & Cohen, 2001). This ability is important to most aspects of human life (Diamond, 2013), as evidenced by the variety of disorders associated with cognitive control dysfunction, such as schizophrenia (Chambon et al., 2008), substance use disorders (Gould, 2010; Wilcox et al., 2014), depression (Marazziti et al., 2010), attention deficit hyperactivity disorder (Barnett et al., 2001), obsessive-compulsive disorder (Abbruzzese et al., 1997), and chronic pain (Berryman et al., 2014; Baker et al., 2016).

The human brain consists of large-scale networks (Power et al., 2011; Yeo et al., 2011). As a higher-order function, cognitive control requires greater coordination among brain-wide networks than simple motor tasks (Cohen & D’Esposito, 2016). Therefore, a prominent hypothesis contends cognitive control arises from dynamic activation of multiple large-scale networks to meet task demands (Menon & D’Esposito, 2022). However, how the brain initiates task-associated network states is not fully understood.

The claustrum is a global network hub, exhibiting the most extensive structural connectivity of all brain regions by volume (Torgerson et al., 2015) and functional connectivity with multiple resting state networks (Krimmel et al., 2019). Synaptic connectivity between the claustrum and the cortex reflects network motifs, and the claustrum is required for cognitively demanding task performance (Qadir et al., 2018; Qadir et al., 2022; White et al., 2018; White & Mathur, 2018a; White & Mathur, 2018b; Atlan et al., 2018; Krimmel et al., 2019; White et al., 2020). To test if the claustrum instantiates multiple networks required for cognitive control (Madden et al., 2022), we analyzed human fMRI data from two executive function tasks evoking different network states and determined whether hypothesized circuits are supported by structural connectivity in corresponding diffusion MRI (dMRI) scans. Claustrum blood oxygenation level dependent (BOLD) activity and circuit analyses were consistent with the claustrum exerting an excitatory influence on cognitive control network nodes across both tasks. These findings support the hypothesis that the claustrum initiates diverse network states that ultimately underpin cognitive control.

## Results

### Claustrum co-activates with externally driven working memory networks

We first sought to test if the claustrum activates networks required for an externally driven task — working memory (Fig. 1A) — in the Amsterdam Open MRI Collection (AOMIC; Snoek et al., 2021). We first observed that trials requiring working memory activated bilateral task positive networks and deactivated task negative networks compared to control trials with no working memory load (Fig 1B). Activity in bilateral claustrum was also greater in working memory trials (Fig 1C). Claustrum BOLD signal was not significantly correlated with motion (Supplemental Table 1).

**Figure 1.**
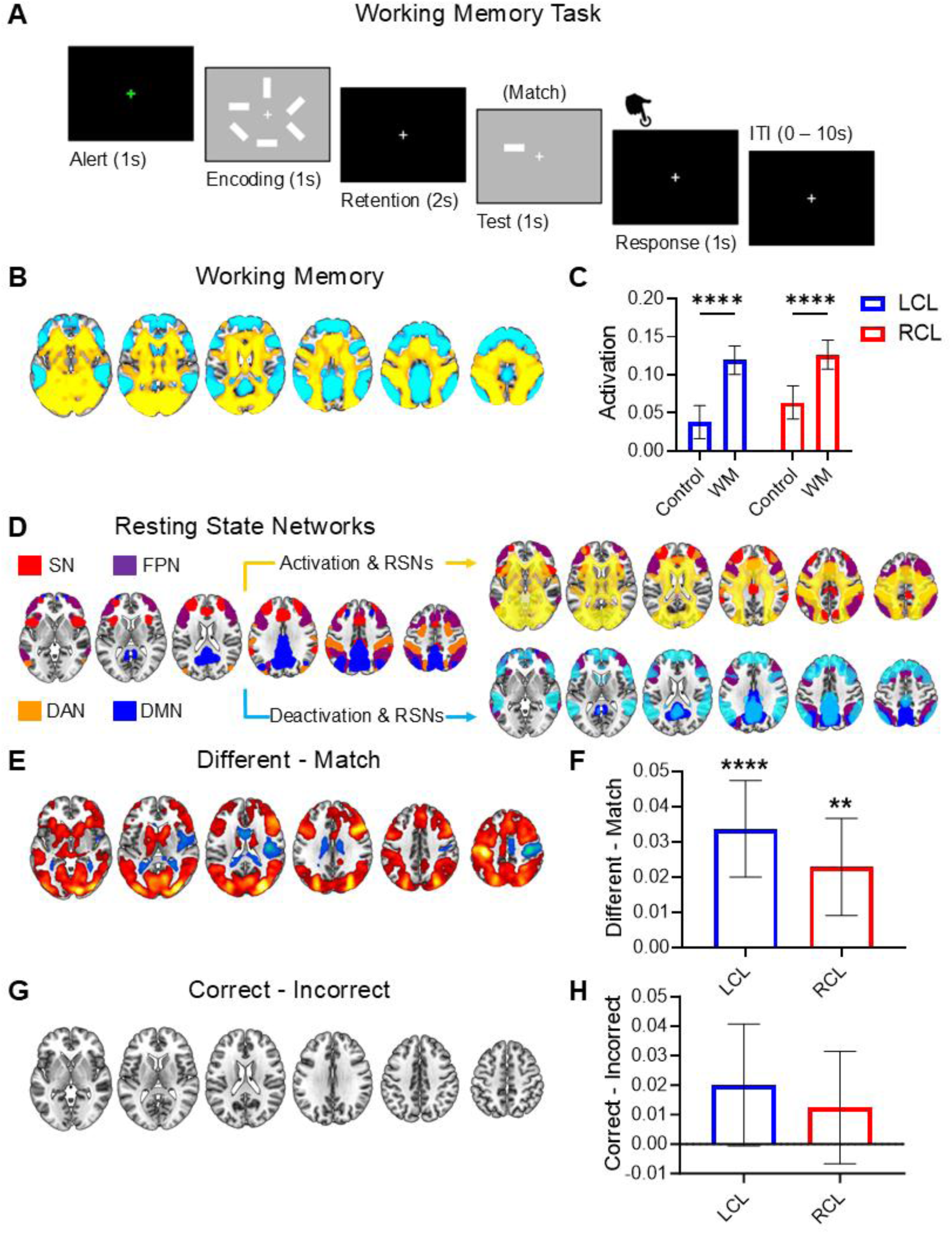
Claustrum co-activates with externally driven working memory networks. **(A)** Working memory scan trials comprised alert, encoding, retention, test, and response phases. **(B)** BOLD signal increases (warm) and decreases (cool) during working memory (*n* = 420). **(C)** Average parameter estimation (regression slope) of condition-related BOLD signal change detected significantly increased bilateral claustrum activation in working memory compared to control trials (two-way ANOVA main effect of condition: *F* (1, 838) = 147.7, *p* < 0.0001; post hoc LCL working memory vs. control: *p* < 0.0001; post hoc RCL working memory vs. control: *p* < 0.0001). No main effect of hemisphere (*F* (1, 838) = 1.536, *p* = 0.2156) or condition x hemisphere interaction (*F* (1, 838) = 2.415, *p* = 0.1206) were detected. **(D)** SN (red), FPN (purple), DAN (orange), and DMN (blue) from resting state group-ICA with working memory BOLD results overlaid. **(E)** “Different - match” trials contrast with greater signal during “different” in red and during “match” in blue. **(F)** Significantly greater LCL (*t* = 4.828, *p* < 0.0001) and RCL (*t* = 3.249, *p* = 0.0013) activity in “different” versus “match” trials. **(G)** No brain-wide differences were observed when contrasting correct and incorrect working memory trials. **(H)** No effects were observed in LCL (Wilcoxon Signed Rank Test *W* = 8928, *p* = 0.0689) or RCL (*W* = 6236, *p* = 0.2039) in “correct - incorrect” contrasts. Whole-brain responses were displayed on horizontal slice montages (z = 0, 10, 20, 30, 40, 50), and results were voxel-wise thresholded at *p* < 0.001, followed by FWE cluster correction. Error bars show 95% confidence intervals. * represents *p* < 0.05, ** < 0.01, *** < 0.001, **** < 0.0001.

Group Independent Component Analysis (ICA) of resting state scans from the same participants identified cognitive control-related networks: the salience network (Seeley et al., 2007; Taylor et al., 2009), fronto-parietal network (Dosenbach et al., 2007; Dosenbach et al., 2008), dorsal attention network (Corbetta & Shulman, 2002; Fox et al., 2006), and default mode network (Greicius et al., 2003) (Fig. 1D). The working memory task was associated with increased activity in salience (e.g., bilateral anterior insula, dorsal anterior cingulate cortex) and dorsal attention (e.g., posterior parietal cortices) network regions, and decreased activity in default mode regions (e.g., posterior cingulate cortex/precuneus) as expected.

During working memory trials, participants judged if a test stimulus differed from or matched the orientation of a rectangle in a previously presented array. Accuracy and reaction times did not differ between “different” and “match” trials (Supplemental Tables 2 & 3). However, the “different – match” contrast revealed differences in whole-brain responses.

Correct responses to “different” trials were made with the right index finger and therefore activated left hemisphere motor cortex. Correct responses to “match” trials were made with the left index finger and so activated right hemisphere motor cortex. Although both trial types evoked bilateral claustrum and network responses (Fig. S3), “different” trials exhibited greater network recruitment compared to “match” trials (Fig. 1E). Bilateral claustrum also exhibited significantly greater activity in the “different - match” contrast (Fig. 1F), illustrating complementary increases in claustrum and network BOLD signal.

When contrasting correct vs. incorrect working memory trials, no differences were detected in whole-brain responses (Fig. 1G) or claustrum activity (Fig. 1H). Therefore, in all analyzed conditions, increased task-associated network signal was always and only observed when claustrum signal increased.

### Claustrum co-activates with internally driven autobiographical memory networks

We next tested claustrum involvement in another task recruiting a different network phenotype: autobiographical memory. Although the default mode network is typically associated with task-induced deactivations, tasks with internally directed demands have demonstrated increased default mode network activity (Spreng et al., 2010; Andrews-Hanna et al., 2014; Adnan et al., 2016; Fuentes-Claramonte et al., 2019) alongside other cognitive control networks. Analysis of a publicly available dataset (*n* = 35; Fuentes-Claramonte et al., 2019) with fMRI scans from an autobiographical memory task (Fig. 2A) confirmed the recruitment of a different network state than working memory, with activations in default mode (e.g., posterior cingulate cortex/precuneus) and salience (e.g., anterior insula, dorsal anterior cingulate cortex) network regions, and deactivations in dorsal attention (e.g., posterior parietal cortex, anterior supramarginal gyrus) network regions (Fig. 2B).

**Figure 2.**
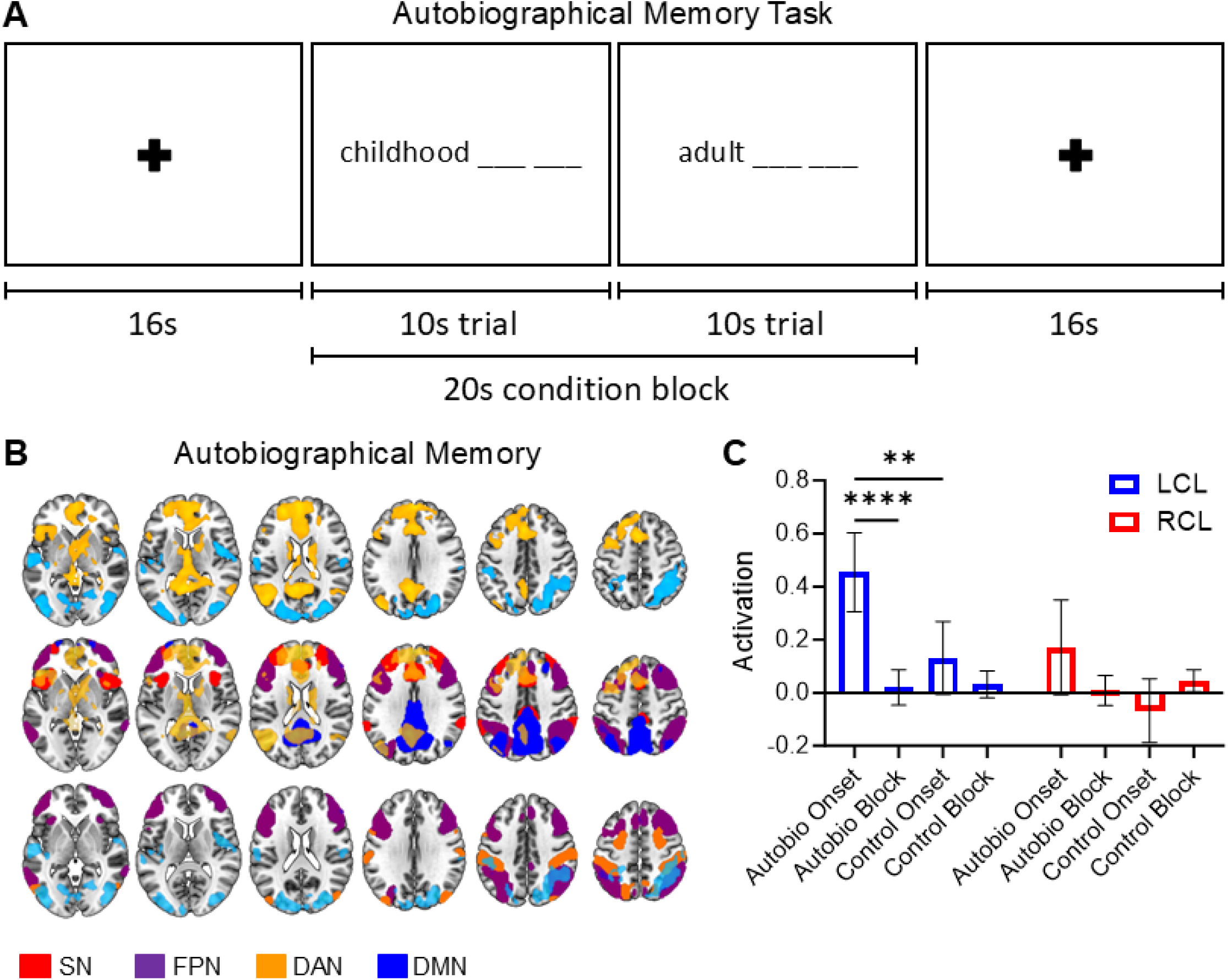
Claustrum co-activates with internally driven autobiographical memory networks. **(A)** Autobiographical memory scans consisted of 20s blocks with 2 consecutive 10s trials. Trials presented a time period label followed by a word pair either corresponding to a participant’s personal memory or not. **(B) Top:** BOLD signal increases (warm) and decreases (cool) during autobiographical memory (*n* = 35) **Middle:** BOLD signal increases from Top overlaid on SN, FPN, and DMN. Note task-related DMN activity (e.g., PCC). **Bottom:** BOLD signal decreases from Top overlaid on FPN and DAN. Displayed RSNs derived from AOMIC data due to absence of resting-state scans in the autobiographical memory dataset. **(C)** LCL, but not RCL, displayed significant activation at autobiographical memory (Autobio) trial onset (two-way ANOVA main effect of condition: *F* (2.129, 144.8) = 13.60, *p* < 0.0001; main effect of hemisphere: *F* (1, 68) = 9.227, *p* = 0.0034; condition x hemisphere interaction: *F* (3, 204) = 3.502, *p* = 0.0164; post hoc LCL Autobio onset vs. Autobio block: *p* < 0.0001; post hoc LCL Autobio onset vs. control onset: *p* = 0.0068).

A significant unilateral activation at the onset of autobiographical memory trials was observed in left claustrum, which was not significantly correlated with subject motion (Supplemental Table 1). Greater left claustrum activation at autobiographical memory trial “onsets” relative to subsequent trial “blocks” and control trial onsets was consistent with cognitive-load dependent claustrum activation at network initiation (Fig. 2C).

Subsequent analyses focused on the left hemisphere, where claustrum activation was observed during working and autobiographical memory. The anterior insula and pulvinar, regions implicated in cognitive control network dynamics (Menon & Uddin, 2010; Molnar-Szakacs & Uddin, 2022; Cortes et al., 2024), also activated during both memory tasks (Figs. S6 and S7). However, left anterior insula activity was less specific to the onset of autobiographical memory conditions than left claustrum (Fig. S8), and left pulvinar activity during autobiographical memory was significantly correlated with subject motion (Supplemental Table 4).

### Claustrum is anatomically connected to cognitive control network nodes

Having observed increased claustrum activity with multiple task network states, we sought to characterize hypothesized claustrum circuits potentially participating in network initiation. We predicted left claustrum would possess structural connectivity with cognitive control network nodes. The anterior cingulate cortex (ACC), premotor cortex (PMC), anterior supramarginal gyrus (SMG), and posterior cingulate cortex/precuneus (PCC) were chosen as representative nodes of the salience, fronto-parietal, dorsal attention, and default mode networks, respectively (Figs. 3A & B). See Methods for node choice rationale.

**Figure 3.**
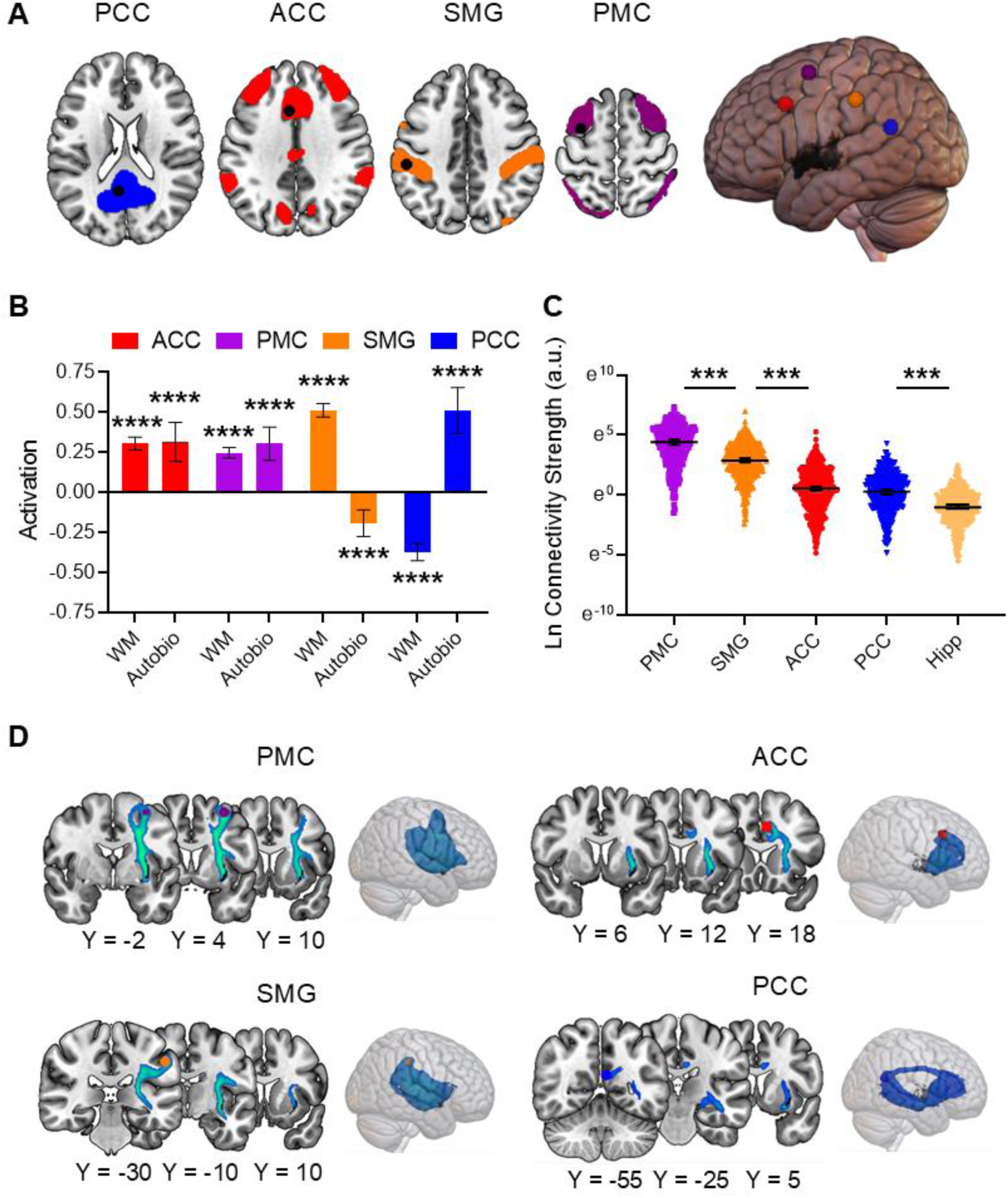
Claustrum is anatomically connected to cognitive task-associated network nodes. **(A)** Horizontal slices (z = 20, 35, 40, 60) individually displaying PCC, ACC, SMG, and PMC ROIs (black circles) on their respective RSNs, and rendered brain with LCL (black). **(B)** Activation parameters in “experimental - control” contrasts in working memory (WM) and Autobio. All regions exhibited contrast values significantly different than zero (ACC WM: *t* = 15.15, *p*-FDR < 0.0001; ACC Autobio: *t* = 5.21, *p*-FDR < 0.0001; PMC WM: *t* = 14.81, *p*-FDR < 0.0001; PMC Autobio: *t* = 5.98, *p*-FDR < 0.0001; SMG WM: *t* = 24.17, *p*-FDR < 0.0001; SMG Autobio: *t* = 4.71, *p*-FDR < 0.0001; PCC WM: *t* = 13.83; *p*-FDR < 0.0001; PCC Autobio: *t* = 7.26, *p*-FDR < 0.0001). **(C)** LCL exhibited preferential structural connectivity with PMC and SMG over ACC and PCC. Select comparisons shown for clarity. **(D)** Group tractograms for LCL connections with network representative nodes thresholded at 50%, meaning fibers detected in at least 50% of participants are shown.

Probabilistic tractography in AOMIC dMRI scans (*n* = 420) revealed significant differences in left claustrum structural connectivity with network nodes (Fig. 3C, χ2(4) = 1068.958, *p* < 0.001). The hippocampus was used as a control, given sparse connectivity between the claustrum and hippocampus (Torgerson et al., 2015; Lei et al., 2025). Post hoc analyses showed that all claustrum connections with network nodes were significantly stronger than the claustrum-hippocampus circuit (Figs. 3C & D; *p*-FWE < 0.001). All structural connectivity *p*-values are shown in Supplemental Tables 5 and 6.

Comparing median structural connectivity strengths (a.u.) of tractograms between claustrum and network nodes revealed preferential claustrum anatomical connectivity with fronto-parietal and dorsal attention network representatives.

The strongest claustrum connectivity was with the PMC (PMC – SMG: *z* = 6.568, *p*-FWE < 0.001), followed by SMG (SMG – ACC: *z* = 12.962, *p*-FWE < 0.001; SMG – PCC: *z* = 14.882, *p*-FWE < 0.001), and finally ACC and PCC, which did not exhibit significantly different claustrum connectivity (ACC – PCC: *z* = 1.92, *p*-FWE = 0.548). Left claustrum also exhibited significantly greater structural connectivity with PMC, SMG, and ACC than the left anterior insula and left pulvinar (Supplemental Table 6).

### Claustrum is driven by higher order association cortices to activate task-related network regions

Given structural data corroborating claustrum connectivity with cognitive control brain networks, we devised a circuit model to investigate how the claustrum is directed by upstream regions to network downstream cortical sites. We tested if any representative network nodes were sources of left claustrum input or targets of left claustrum output. Because ACC input to the claustrum is necessary for optimal performance in a cognitively demanding task in mice (White et al., 2018) and the claustrum robustly relays ACC input to frontal and posterior cortical regions (Qadir et al., 2022), we predicted that ACC would supply driving input to the claustrum, which would exert excitatory influence on task-related nodes.

Dynamic causal models (DCM; Friston et al., 2003) were defined ranging from a full model in which memory tasks modulated all claustrum projections to a null model in which no claustrum projections were task-responsive (Figs. 4A & B). This set of models was crossed with five possible sources of claustrum input, including the four representative nodes of networks of interest: ACC (salience), PMC (fronto-parietal), SMG (dorsal attention), PCC (default mode); and the claustrum itself, which represented all other potential input sources.

**Figure 4.**
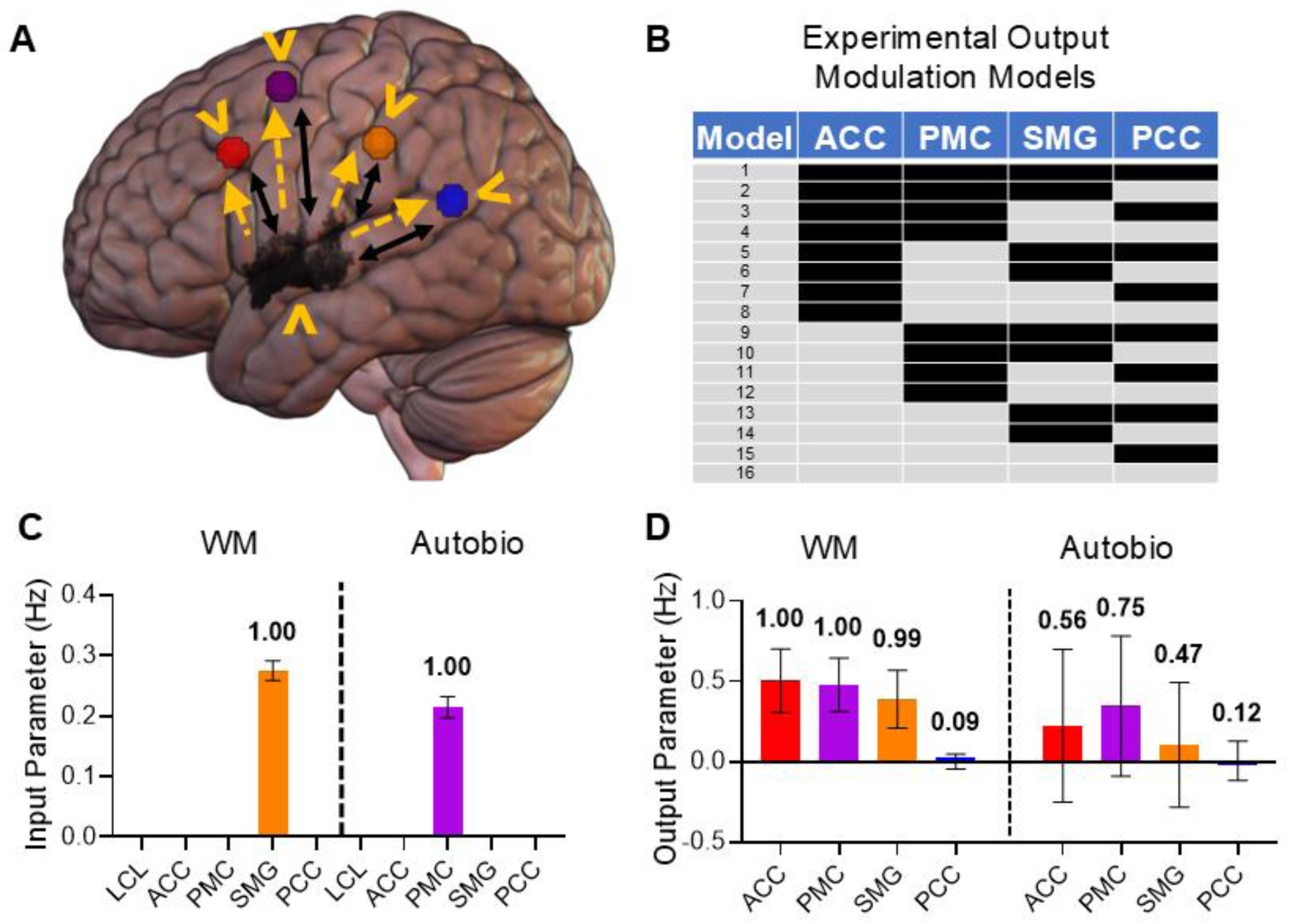
Claustrum is driven by higher order association cortices to activate task-related network regions. **(A)** The “full” DCM contained all verified anatomical connections (black arrows), possible driving experimental input (orange ‘^’ symbols) from each node, and possible task-induced modulation of all LCL outputs (orange dashed arrows). Tested models represented all 16 possible combinations of experimental claustrum output modulation from a full model (#1) to a null model (#16). **(C) Left:** Estimated input parameters during WM and Autobio **(D)** Estimated output parameters during WM and Autobio. 95% confidence intervals and posterior probabilities signifying evidence strength (> 0.5 = “weak”, > 0.75 = “positive”, > 0.95 = “strong”, > 0.99 = “very strong”) accompany bars.

Positive input parameters and high posterior probabilities indicated very strong evidence for SMG input to left claustrum during working memory and PMC input during autobiographical memory (Fig. 4C). Comparisons between tasks revealed very strong evidence for greater SMG input to claustrum during working memory (Fig. S9).

Output parameters reflected strong or very strong evidence for excitatory left claustrum effective connectivity to ACC, PMC, and SMG, but not PCC during working memory, as well as weak evidence for excitatory left claustrum influence on ACC and PMC during autobiographical memory (Fig. 4D). Comparisons between tasks revealed weak evidence for greater left claustrum excitatory influence on ACC and SMG in working memory (Fig. S9). Posterior probabilities for left claustrum were consistently stronger than left anterior insula and left pulvinar when assessing excitatory effective connectivity with ACC, PMC, and SMG (Figs. S6 and S7). All Bayesian output model comparison probabilities are shown in Supplemental Table 7.

A reciprocal relationship was observed between left claustrum and SMG, a dorsal attention network node, during the externally driven working memory task. PMC, a fronto-parietal network node, was identified as a source of input to claustrum during autobiographical memory and a task-general recipient of claustrum output. The ACC, a salience network node, was observed to be excited by left claustrum during both tasks, with stronger evidence of this during working memory. No analyses supported left claustrum input from or output to PCC.

These findings align with observed preferential claustrum structural connectivity with SMG and PMC, indicating the claustrum is driven by input from higher order association cortices to excite task-related regions in fronto-parietal, dorsal attention, and salience, but not default mode, networks.

## Discussion

Our results implicate the claustrum in initiating multiple networks supporting diverse cognitive demands. Circuit analyses suggest that the claustrum excites a task-related subset of its anatomical connections in a cognitive load-dependent manner, and that the claustrum shares stronger anatomical and functional connections with fronto-parietal and dorsal attention than salience and default mode network nodes. Greater evidence for a role of the claustrum in network activation than pulvinar and anterior insula establish this long-understudied structure as a key player in network dynamics underlying cognitive control.

### Claustrum effects coincide with cognitive control network changes

Working memory evoked similar networks if the test stimulus matched the remembered stimuli or not, but network BOLD signal intensity was greater when the test and remembered stimuli differed. Analogously, claustrum activity was observed in both conditions, and claustrum activity was significantly greater in “different” than “match” trials. Conversely, no differences were detected in brain-wide networks or claustrum signal between correct and incorrect trials. Therefore, network effects always and only appeared alongside claustrum effects.

Working memory test condition effects on network or claustrum BOLD signal must have arisen while processing the test stimulus, executing the motor response, or both. “Different” and “match” trials required button presses from opposite hands, which was evident in opposing unilateral motor cortex responses, but increased claustrum activity was bilateral in both conditions. Network and claustrum BOLD effects likely also occurred in earlier task phases, such as encoding and retention, a claim supported by similar patterns of network activity and increased bilateral claustrum signal when working memory trials were compared to control trials, which included motor responses. Left claustrum signal at autobiographical memory trial onset, during a task with no motor response, is further evidence that claustrum supports initiation of cognitive control networks. Findings in both memory tasks, which evoked starkly different network motifs, implicate the claustrum in the appearance of cognitive control networks across a wide range of network states.

### Structural and functional claustrum-network relationships

dMRI analyses revealed tiers of claustrum structural connectivity. Claustrum-hippocampus connectivity was the weakest analyzed, consistent with findings in macaques and humans motivating its use as a control (Torgerson et al., 2015; Lei et al., 2025). Claustrum connections with ACC and PCC, representative nodes of salience and default mode networks, were significantly stronger than claustrum-hippocampus, and claustrum connections with SMG and PMC, representatives of dorsal attention and fronto-parietal networks, were the strongest observed. This did not match our expectation that ACC would possess the strongest claustrum connectivity given the importance of claustrum-ACC connections observed in other species (White et al., 2017; Qadir et al., 2018; White & Mathur, 2018a; White & Mathur, 2018b; White et al., 2018; Xu et al., 2022; Ntamati et al., 2023; Faig et al., 2024). It is possible this resulted from measurement error. dMRI tractography connection weights correlate with those derived from tracer injections, but the method is not perfect (Donahue et al., 2016). However, these findings may also reflect species differences due to significant human cortical expansion (Preuss & Wise, 2022).

Preferential claustrum connectivity with dorsal attention and fronto-parietal networks is supported by a previous claustrum dMRI analysis (Stewart et al., 2024) and by this study’s DCM, which consistently favored SMG and PMC as sources of task-relevant input to claustrum over other network nodes. DCM also found evidence for excitatory claustrum projections to PMC in the fronto-parietal network during both tasks and to SMG during working memory, which recruited the dorsal attention network. DCM supported excitatory claustrum projections to ACC in both tasks as well, but the absence of claustrum-PCC effective connectivity is conspicuous and reinforces the general agreement between DCM and dMRI, which found relatively weak claustrum-PCC structural connectivity.

The effect of claustrum projections on their targets varied based on several factors (McBride et al., 2023), and we have speculated previously on the possibility of claustrum interactions with the default mode network (Madden et al., 2022). While we predicted claustrum supported network initiation, we were uncertain if that entailed excitation of task-positive regions, inhibition of task-negative regions, or both. DCM suggested that the claustrum exerts task-dependent excitation on ACC, PMC, and SMG. No evidence of claustrum influence on PCC was found, so the functional relevance of the observed claustrum-PCC structural connection may apply in other brain states, such as sleep (Marriott et al., 2024).

DCM analyses therefore reflected circuits where task-related association cortices drive the claustrum to support network activity among a task-related subset of its anatomical connections.

### Claustrum role in cognitive control network initiation

Control analyses of the pulvinar and anterior insula revealed different results than the claustrum. The pulvinar nuclei connect with prefrontal and posterior parietal cortices associated with cognitive control networks (Yeterian & Pandya, 1985; Romanski et al., 1997; Leh et al., 2008), and task-based fMRI has implicated the region in working memory (Rotshtein et al., 2011). Consistent with this literature, we observed increased medial pulvinar BOLD signal during working memory, but evidence for pulvinar activity during autobiographical memory was less robust. A significant BOLD signal increase at autobiographical memory trial onset may have been artifactual as it shared a significant positive correlation with subject movement. Moreover, DCM evidence for excitatory pulvinar output during the task was comparable to that of the null model, which contained no task-induced modulation of pulvinar projections. Lastly, pulvinar exhibited preferential structural connectivity with the hippocampus, the negative control region for claustrum structural connectivity analyses. These findings are consistent with a pulvinar role in cognitive control network coordination, but the case is strongest in the externally driven working memory task.

The anterior insula shares connections with cognitive control associated cortical regions (Mesulam & Mufson, 1982; Mufson & Mesulam, 1982) and responds to a wide variety of stimuli prior to other brain regions (Sridharan et al., 2008). Experimental support has even been found for a causal influence of anterior insula activity on cognitive control network regions; however, this evidence has been shown in relatively simple tasks (Sridharan et al., 2008). Analyses of anterior insula effective connectivity specific to conditions of increased cognitive demand find causal influence restricted to the dorsal anterior cingulate, the anterior insula’s salience network counterpart (Cai et al., 2016). This literature is consistent with anterior insula recognizing cognitive control needs but downstream mechanisms contributing to demand-dependent network activation. We observed significant anterior insula BOLD increases during both memory tasks.

Notably, anterior insula responses during autobiographical memory scans extended into trial “blocks” and were detected in response to control stimuli, whereas claustrum effects were observed specifically at the *onset* of autobiographical memory trials. Surprisingly, DCM found strong evidence for inhibitory influence of anterior insula outputs on network representative nodes. Given contrary findings in prior studies using different effective connectivity techniques, the limitations of DCM prevent us from concluding anterior insula projections are inhibitory. Nevertheless, opposing anterior insula and claustrum DCM results, distinct task response profiles, and superior claustrum structural connectivity with network representative nodes are consistent with the regions performing different roles in network mechanisms of cognitive control.

### Limitations

This study’s findings are subject to important limitations. Bilateral claustrum responses were observed during working memory, but only left claustrum responses were significant during autobiographical memory. This may be due to differences in sample size and handedness between the datasets, but task differences in claustrum effects could have arisen from laterality in claustrum function, a possibility evinced in other human claustrum studies (Barrett et al., 2020; Stewart et al., 2024). Our focus on task differences restricted circuit analyses to the left hemisphere, where claustrum effects were observed in both datasets, leaving laterality in structural connectivity and effective connectivity unexplored. Our circuit analyses could not reveal the relative contributions of afferent and efferent projections to claustrum structural connectivity because dMRI is agnostic to the direction of fibers. Finally, although other connections and nodes are undoubtedly involved in mechanisms of cognitive control network activity, we opted to focus effective connectivity model complexity on circuit aspects relevant to this study’s research questions: inputs to, and outputs from, the claustrum.

### Conclusion

Significant claustrum responses were observed to externally and internally driven cognitive control demands, which evoked markedly different network states. These findings converged with structural and effective connectivity analyses to implicate the claustrum in supporting activation of multiple, large-scale cognitive control networks. Control analyses revealed distinct task response, anatomical connectivity, and effective connectivity profiles for the claustrum, anterior insula, and pulvinar, suggesting complementary roles for these regions in directing network dynamics. The claustrum exhibited the strongest structural connectivity with cognitive control network nodes and displayed excitatory effective connectivity with recruited nodes across tasks. This evidence suggests that the claustrum supports the initiation of cognitive control network states for externally and internally driven cognitive control demands.

## Materials and Methods

### Participants

Analyses were performed in three datasets. The first two datasets are components of the publicly available, multimodal Amsterdam Open MRI Collection (AOMIC; https://nilab-uva.github.io/AOMIC.github.io/). Full details on participant inclusion and demographic variables can be found in Snoek et al. (2021). Briefly, datasets labeled PIOP1 and PIOP2 consisted of university students at the Amsterdam University of Applied Sciences and the University of Amsterdam. Of the manifold AOMIC scans available, this study analyzed working memory task, resting-state, and dMRI scans. Because structural and effective connectivity analysis ROIs were selected based on task responses and resting state network membership, all analyses were performed on the subset of participants who possessed all three scan types (PIOP1: *n* = 198, 108 female, 83 male, 7 unspecified, mean +/- sd. age = 22.20 +/- 1.81 years; PIOP2: *n* = 222, 126 female, 95 male, 1 unspecified, mean +/- sd. age = 21.95 +/- 1.79 years). Of the 216 PIOP1 participants, 18 were excluded due to the absence of a desired scan. Of the 226 PIOP2 participants, 4 were excluded due to the absence of a desired scan. When initial analyses of PIOP1 were reproduced in PIOP2 (Fig. S1-S5), the datasets were combined. All main text results were derived from the pooled sample (n = 420).

The third dataset was obtained by Fuentes-Claramonte et al. (2019) and was selected for this study for analysis of an autobiographical memory task (https://openneuro.org/datasets/ds001618/versions/1.0.1). After the exclusion of 1 participant due to excessive motion, this dataset yielded *n* = 35 (mean +/- sd. age = 40.89 +/- 12.02 years, all right-handed). The authors of both studies obtained approval from their overseeing ethics committees prior to data acquisition, and all participants provided written informed consent.

### Imaging Acquisition and Preprocessing

#### AOMIC

Full details of scan acquisition can be found in Snoek et al. (2021). Both PIOP1 and PIOP2 datasets were acquired on the same Phillips 3T scanner (Philips, Best, the Netherlands) with a 32-channel head coil. PIOP1 was acquired on the “Achieva” version, and PIOP2 was acquired later, after the scanner was upgraded to the “Achieva dStream” version. Both datasets acquired high-resolution 3D MPRAGE anatomical scans (repetition time/echo time [TR/TE] 8.5/3.9ms, slice thickness 1mm, field of view [FOV] 188 × 240 × 220mm, flip angle 8°, voxel size 1 × 1 × 1mm).

All AOMIC data was standardized in Brain Imaging Data Structure format and preprocessed with Fmriprep version 1.4.1 (RRID:SCR_016216; Esteban et al., 2019; Esteban et al., 2020), a Nipype based tool (RRID:SCR_002502; Gorgolewski et al., 2011; Gorgolewski et al., 2017).

#### Working Memory

PIOP1 and PIOP2 working memory functional data were acquired with a gradient echo echo-planar imaging (EPI) sequence (TR = 2000ms, TE = 28ms, 37 slices, FOV = 240 x 240 x 122mm, flip angle = 76.1°, voxel size 3 x 3 x 3mm, slice gap = 0.3mm).

Publicly available preprocessed functional images were unsmoothed. Additional preprocessing was performed in SPM12 (https://www.fil.ion.ucl.ac.uk/spm/software/spm12/). Images were resampled to 2 x 2 x 2mm voxels with a 4th degree B-Spline interpolation. Claustrum analyses used unsmoothed images to prevent inclusion of extra-claustral signal in claustrum ROI voxels. All other analyses used images smoothed with a 6mm full width at half maximum (FWHM) Gaussian kernel.

#### Resting-State

PIOP1 resting state data were acquired with a gradient echo EPI sequence (TR = 750ms, TE = 28ms, 36 slices, FOV = 240 x 240 x 118mm, flip angle = 60°, voxel size 3 x 3 x 3mm, slice gap = 0.3mm). PIOP2 resting state data were acquired with a gradient echo EPI sequence (TR = 2000ms, TE = 28ms, 37 slices, FOV = 240 x 240 x 122mm, flip angle = 76.1°, voxel size 3 x 3 x 3mm, slice gap = 0.3mm). PIOP1 resting state scans lasted 6 minutes, and PIOP2 resting state scans lasted 8 minutes. PIOP1 resting state scans were acquired with multi-slice acceleration (factor 3), while PIOP2 scans were not, resulting in different TRs.

Publicly available preprocessed functional images were unsmoothed. Additional preprocessing was performed in in SPM12 (https://www.fil.ion.ucl.ac.uk/spm/software/spm12/). Images were resampled to 2 x 2 x 2mm voxels with a 4th degree B-Spline interpolation and smoothed with a 6mm FWHM Gaussian kernel.

Participants were directed to maintain their gaze on a fixation plus sign in the middle of a screen and to let their thoughts run freely.

#### Diffusion Weighted Imaging

PIOP1 and PIOP2 used spin-echo diffusion imaging over a single run (TR = 7387ms, TE = 86ms, slice thickness = 2mm, FOV = 224 × 224 × 120mm). Diffusion weighting consisted of a single shell (b = 1000s/mm^2^). Each scan contained 32 diffusion-weighted directions, and a single non-diffusion weighting image (b0).

Preprocessed dMRI scans were downloaded from the AOMIC database, which underwent a custom preprocessing pipeline utilizing tools from both FSL (the FMRIB Software Library; Jenkinson et al., 2012) and MRtrix3 (Tournier et al., 2019). MRtrix3 was used for denoising (dwidenoise; Veraart et al., 2016a; Veraart et al., 2016b), Gibbs artifact removal (mrdegibbs; Kellner et al., 2016), eddy current and motion correction (dwipreproc), which utilizes FSL’s eddy (Andersson & Sotiropoulos, 2016), bias correction (dwibiascorrect; Tournier et al., 2019), brain mask extraction (dwi2mask; Dhollander & Connelly, 2016), and gradient diffusion correction (dwigradcheck; Jeurissen et al., 2014). After preprocessing, a diffusion tensor model was fit to the data (dwi2tensor; Veraart et al., 2013), a fractional anisotropy image was extracted (tensor2metric; Basser et al., 1994), and finally a population fractional anisotropy template was created (population_template).

#### Autobiographical Memory

Full details of scan acquisition can be found in Fuentes-Claramonte et al. (2019). Autobiographical memory scans were acquired with a 3T Philips Achieva scanner (Philips Medical Systems, Best, The Netherlands). Functional data were acquired with a T2*-weighted EPI sequence (TR = 2000ms, TE = 30ms, slice thickness = 3mm, FOV = 240mm, flip angle = 78°, in-plane resolution = 3 × 3mm, inter-slice gap = 1mm). Thirty-two slices were acquired per volume with an ascending order parallel to the AC-PC plane. Fuentes-Claramonte et al. (2019) discarded the first 10 volumes to avoid T1 saturation effects. In this study, all volumes were included because no differences were noted in claustrum signal when discarding or including initial volumes. If anything, claustrum response values marginally increased when discarding initial volumes, potentially biasing results toward significant effects (Fig. S10). High-resolution anatomical scans were acquired but were not shared publicly (https://openneuro.org/datasets/ds001618/versions/1.0.1) and were therefore not used in image preprocessing.

All autobiographical memory scans were preprocessed in SPM12 (https://www.fil.ion.ucl.ac.uk/spm/software/spm12/). Preprocessing included realignment (motion correction) and coregistration and normalization of realigned functional images to SPM’s template mean EPI image file (Calhoun et al., 2017) in Montreal Neurological Institute (MNI) space with interpolation to 2 x 2 x 2mm voxels using a 2nd degree B-Spline. Claustrum analyses used unsmoothed images to prevent inclusion of extra-claustral signal in claustrum ROI voxels. Whole-brain task response analyses used images smoothed with a 6mm FWHM Gaussian kernel.

### Tasks

#### Working Memory

The working memory task (Fig. 1A) was designed to measure brain processes related to visual working memory (Snoek et al., 2021) using a protocol inspired by Pessoa et al. (2002). Each run consisted of 40 trials: 32 trials which possessed a working memory load, and 8 trials which did not, but were designed to control for visual stimulation and motor responses.

During the inter-trial interval (ITI), participants attended to a white fixation plus sign on a black background. Trials began with a change of the plus sign’s color from white to green, which lasted for 1s, alerting the participant to trial onset. In working memory trials, this was followed by a change in background color from black to gray and the presentation of an array of 6 white bars arranged in a circle with random orientations, which appeared for 1s. Control trials only included the change in background color. After this, the white fixation plus sign was displayed on the original black background for 2s. Subsequently, working memory trials displayed a gray background with a test stimulus of one bar. After 1s of test stimulus presentation, the background returned to black, and participants had 1s to respond if the test stimulus matched (16 trials, left index finger press) or differed from (16 trials, right index finger press) a bar in the trial’s previously displayed array.

Instead of displaying a test stimulus, control trials similarly changed background from black to gray, but displayed the direction to “respond left” or “respond right” for 1s, which the participant was instructed to perform during the 1s response phase after the direction disappeared and the background returned to black.

### Autobiographical Memory

Immediately prior to the fMRI session, a member of the data-acquiring research team (Fuentes-Claramonte et al., 2019) interviewed each participant for approximately 1 hour. The interview consisted of prompts from the Autobiographical Memory Interview (AMI; Kopelman et al., 1989) and the Crovitz test (Crovitz & Schiffman, 1974) to identify four to six memories from each of four time periods: “childhood” (before age 12), “adolescence” (ages 12–18), “adulthood” (after age 18) and “recently” (within the last year). Autobiographical memory stimuli were three-word prompts, comprising a time period label and a word pair signifying a previously identified memory. Word pairs were decided collaboratively during the interview by the researcher and participant, and word pairs were only generated for memories receiving a maximum AMI score of 3 for the memory’s level of detail and specificity in time and place. Control stimuli, designed to align with autobiographical memory stimuli in terms of perceptual and linguistic processing (Fuentes-Claramonte et al., 2019), were also three-word prompts, but comprising a time period label followed by a word pair with no reported memory association. Scans consisted of 20 stimulation blocks separated by 16s long ITIs where a fixation plus sign was presented (Fig. 2A).

Blocks alternated between control stimuli and autobiographical stimuli, and each 20s block consisted of 2 trials, each lasting 10s. Participants were directed to read the words presented at each trial silently and recall any associated memory.

### Task Modeling

Previous analyses of task-induced claustrum BOLD signal change have used hybrid event-block designs (Krimmel et al., 2019; Stewart et al., 2024) where single condition blocks were modeled with two regressors, one representing a brief condition “onset” period and another representing the remainder of the condition “block”. This modeling format has tested the prediction that transient claustrum activation occurs at task onset to support task-associated network initiation, and claustrum BOLD signal increases have been observed specifically at the onset of a difficult cognitive task condition (Krimmel et al., 2019) and the onset of acute pain (Stewart et al., 2024).

### Working Memory

Although models representing each component of the working memory task (i.e., alert, encoding, retention, test, response) were desired, the brief duration (1-2s) of the trial components resulted in highly correlated regressors and a high likelihood of effect misattribution (Supplemental Table 8). Therefore, working memory scan GLMs contained three regressors, representing the entire duration of each trial type (“working memory different,” “working memory match,” “control”), capturing ITI in the implicit baseline. For DCM analyses (see below), a second model with two regressors (“experimental,” “control”) was used where “working memory different” and “working memory match” were pooled into one “experimental” condition. Results from this model are presented in Fig. 1B.

### Autobiographical Memory

Low VIFs confirmed hybrid event-block GLMs were appropriate for the autobiographical memory task (O’Brien, 2007; Supplemental Table 9). Therefore each 10s trial (2 per block) was modeled separately as a 2s “onset” condition beginning at the trial onset followed by an 8s “block” condition lasting the remainder of the trial. Separate “onset” and “block” regressors were used for autobiographical memory and control trials. As in the working memory task, the ITI was captured in the implicit baseline. For DCM analyses (see below), a second model with two regressors (“experimental,” “control”) was used where “onset” and “block” were combined yielding 10s regressors lasting the duration of each trial.

All task fMRI analyses were performed in SPM12. For analyses of ROIs other than the claustrum, GLMs included all task conditions and six motion parameters. Second level results were separately masked with each ROI and activation values were averaged across the ROI within each subject, and then across subjects. Separate GLMs were used to analyze each hemisphere’s claustrum ROI, which included all task conditions, six motion parameters, and the two flanking region-condition interaction timeseries generated via the task-adapted SRCC process described below.

### Small Region Confound Correction

As described previously (Krimmel et al., 2019), the effects of neighboring insular cortex and putamen on claustrum signal were controlled via Small Region Confound Correction (SRCC). Insular cortex and putamen “flanking” ROIs were defined by dilating each hemisphere’s claustrum ROI four functional voxels and identifying the overlap between the dilated claustrum and the neighboring insula and putamen at least two functional voxels separated from the original claustrum. This generated “flanking” ROIs within the insular cortex and putamen, similar to the claustrum’s shape, yet apart from the claustrum to avoid including claustrum signal.

In previous claustrum resting state connectivity analyses (Krimmel et al., 2019), the effects of flanking regions were controlled by including their time series as additional regressors. However, if flanking ROI signals are influenced by task conditions, using physiological flanking ROI time series as additional regressors in task-based claustrum analyses risks removing condition-induced variation from the claustrum signal (Stewart et al., 2024). Therefore, when analyzing claustrum task responses, the CONN Toolbox (RRID:SCR_009550; Nieto-Castanon, 2020) was used to generate canonical hemodynamic response function (HRF) convolved condition time series, and a regressor was generated for each flanking ROI by obtaining the de-meaned element-wise product of the ROI’s de-meaned physiological time series and the de-meaned, summed HRF-convolved time series of all modeled conditions. This yielded a time series for each flanking ROI that covaried with the ROI physiological time series during task conditions but lacked variation potentially induced by the conditions, allowing control of the influence of neighboring regions on claustrum signal without indirectly removing task effects.

### Resting State Group-ICA

Cognitive control RSNs were derived from PIOP1 and PIOP2 preprocessed, smoothed resting state scans in the CONN Toolbox (RRID:SCR_009550; Nieto-Castanon, 2020). Nuisance regressors included motion parameters and their first order derivatives, a scrubbing vector generated by ART-toolbox identification of outlier scans (global-signal z-value threshold = 5; subject-motion mm threshold = 0.9), the first five principal components of white matter and cerebrospinal fluid (CSF) masks (aCompCor; Behzadi et al., 2007; Muschelli et al., 2014), and a default regressor in the CONN Toolbox to control for potential ramping effects at scan onset. Global signal regression was not performed. Consequently, a 2x eroded CSF mask and a 4x eroded white matter mask were used because such masks no longer contain global signal (Power et al., 2017). Regression was performed simultaneously with band-pass filtering (Hallquist et al., 2013) between 0.008Hz and 0.09Hz, and linear detrending was also performed.

Group-ICA was performed in the CONN Toolbox following methods described in Calhoun et al. (2001) using individual subject dimensionality reduction to 64 dimensions prior to G1 FastICA and GICA3 Back-projection. RSNs were selected from resulting spatial components based on comparisons with relevant literature (Corbetta & Shulman, 2002; Greicius et al., 2003; Fox et al., 2006; Dosenbach et al., 2007; Seeley et al., 2007; Dosenbach et al., 2008; Taylor et al., 2009; Witt et al., 2021) and the CONN Toolbox’s “spatial match to template” tool, which calculates component correlations with CONN’s default network atlas, derived from a group-ICA of Human Connectome Project data (*n* = 497). PIOP1 was subjected to a 10 component parcellation, which yielded all desired networks (salience, fronto-parietal, dorsal attention, default mode). However, a 10-component ICA did not robustly capture all desired networks in PIOP2 or when combining the datasets. Therefore, a 15-component ICA was used to define networks in PIOP2 specifically and when combining the datasets.

### Regions of Interest

All ROIs are publicly available on NeuroVault (https://neurovault.org/collections/16667/).

LCL, RCL, insular cortex, and putamen ROIs were drawn by hand on the normalized anatomical images of 20 subjects in a publicly available dataset (*n* = 22) acquired on a 7T MR Scanner (MAGNETOM 7T, Siemens Healthcare, Erlangen, Germany), and a group average ROI file was obtained for each region. Further details of the scans can be found in Gorgolewski et al (2015). Two subjects were omitted from ROI generation, one due to preprocessing errors preventing normalization of sufficient quality and one due to acquisition via different scanning parameters. The relevance of insular cortex and putamen ROIs, as well as a description of “flanking” ROIs can be found in the “Small Region Confound Correction” Methods above. All SRCC-related ROIs are publicly available.

Circuit analysis ROIs were selected based on the overlap of “experimental - control” trial contrast maps (working memory: = experimental, control = control; autobiographical memory: autobiographical = experimental, control = control) and group-ICA resting state network maps. All ROIs were in the left hemisphere because only LCL exhibited significant responses to both memory tasks, and because recent tracing experiments in macaques observed a strong ipsilateral bias for both claustrum afferents and efferents (Lei et al., 2025), a finding supported by recent tractography analyses in humans (Wendt et al., 2024).

Salience network activity appears in response to a wide variety of tasks and has been observed to temporally precede activity in other network regions (Sridharan et al., 2008). It has therefore been proposed to “gate” the switching between default mode and task-positive network states (Menon & Uddin, 2010; Molnar-Szakacs & Uddin, 2022). ACC is a primary salience network node, and increasing evidence attests to the functional importance of bidirectional ACC-claustrum projections (White & Mathur, 2018a; White & Mathur, 2018b; White et al., 2018). We therefore selected a salience network ROI in ACC (center: −6, 18, 36) that was active in both memory tasks.

A network of brain regions consistently active across a wide variety of cognitive control tasks has been referred to as the multiple demand network (Duncan, 2010; Fedorenko et al., 2013), and this network has been shown to consist primarily of fronto-parietal network regions (Assem et al., 2020). We therefore selected an ROI active in both tasks that was centered within the data-derived fronto-parietal network and a publicly available multiple demand network map (https://imaging.mrc-cbu.cam.ac.uk/imaging/MDsystem). This ROI resided in PMC (center: −32, 2, 58).

The posterior cingulate cortex/precuneus (PCC) is a hallmark default mode network node (Greicius et al., 2003), and the memory tasks were selected due to their predicted opposing effects on DMN activity. Therefore, a PCC ROI (center: −6, −54, 20) within the data-derived default mode network that exhibited differential activation between memory tasks was selected. Conversely, the dorsal attention network is characterized by its anticorrelation with the default mode network (Fox et al., 2005; Fox et al., 2006). Therefore, an ROI from the AOMIC-derived dorsal attention network exhibiting a task response profile opposing that of PCC was selected. This ROI resided in SMG (center: −54, −30, 40).

Circuit analysis ROIs overlaid on more complete views of the resting state networks than depicted in Fig. 3A can be found in Figure S11.

Left anterior insula location (center: −38, 18, −4) was selected in an area responsive to both memory tasks, similarly to ACC, as a salience network representative.

To define the left pulvinar ROI, a publicly available functional pulvinar parcellation (Guedj & Vuilleumier, 2020) derived from a previous anatomical parcellation (Krauth et al., 2010) was analyzed for memory task responses. The working memory task evoked relatively uniform responses across pulvinar parcels, whereas the autobiographical memory task evoked signal increases from ventromedial and dorsomedial parcels. These two parcels were combined and masked with SPM12’s cerebrospinal fluid tissue map due to the region’s proximity to ventricles. Human tractography confirms significant ipsilateral pulvinar structural connectivity (Leh et al., 2008).

For LCL structural connectivity analyses, a negative control region was selected in the left hippocampus (Hipp; center: −26, −20, −16) based on evidence of weak claustrum-hippocampus anatomical connectivity (Torgerson et al., 2015; Lei et al., 2025).

All circuit analysis ROIs except left pulvinar were generated in MarsBaR software (Brett et al., 2002) as 5mm radius spheres.

### Structural Connectivity

AOMIC dMRI scans underwent fiber tract estimation and orientation using FSL’s Bayesian Estimation of Diffusion Parameters Obtained using Sampling Techniques-Crossing Fibers (BEDPOSTx; Behrens et al., 2003; Behrens et al., 2007).

Probabilistic tractography was run using FSL’s Probtrackx2 GPU version (Behrens et al., 2003; Behrens et al., 2007; Hernandez-Fernandez et al., 2019) to assess structural connectivity of LCL, LaINS, and LPulv with ACC, PMC, SMG, PCC, and Hipp. Each ROI was in MNI152 space and was non-linearly transformed to each individual’s diffusion space using FMRIB’s Non-linear Image Registration Tool (FNIRT; Jenkinson et al., 2012), during Probtrackx2 analysis. For each connection (e.g., LCL-ACC), two tractograms were computed, one for each direction, to control for directional biases in dMRI acquisition (Van Essen et al., 2013) and fiber fanning (Jeurissen et al., 2019). The modified Euler algorithm was used to generate the tracts with 10,000 streamlines per voxel. To exclude spurious connections, we included exclusion masks that guided tractography. All tractogram analyses used the following exclusion masks:

i. a combination of ACC, PMC, SMG, PCC, and Hipp masks depending on the analyzed target ROI (e.g., when assessing LCL-ACC connectivity, PMC, SMG, PCC, and Hipp exclusion masks were used),
ii. a combination of LCL, thalamus (Frazier et al., 2005; Desikan et al., 2006; Makris et al., 2006; Goldstein et al., 2007), and insular cortex (Cormie et al., 2023) masks, depending on the analyzed seed ROI (i.e., LCL analyses used the insular cortex and thalamus masks, LaINS analyses used LCL and thalamus masks, and LPulv analyses used LCL and insular cortex masks),
iii. an amygdala mask (Frazier et al., 2005; Desikan et al., 2006; Makris et al., 2006; Goldstein et al., 2007),
iv. a mid-sagittal mask, to remove tracts crossing the midline, and
v. target-specific coronal and/or transverse plane masks to further constrain connections and avoid spurious tracts (ACC: z = 52, y = 50, y = −60; PMC: y = −52; SMG: y = −58; PCC: z = 42, y = −68; Hipp: z = 18, z = −32).

An example structural connectivity exclusion mask setup is illustrated in Figure S12.

To control for streamline propagation, we configured each tractography analysis to propagate toward and not past the target ROI. This means for probabilistic tractography analyses, the target ROI was used as both the termination and waypoint mask. For example, when analyzing the connection from LCL to ACC, ACC was used as the termination and waypoint mask.

Structural connectivity strength was operationalized as the number of streamlines that arrived at the target from the seed region (‘waytotal’). As Probtrackx2 was set to send 10,000 streamlines from each voxel of the seed mask, the total number of streamlines sent is 10,000 multiplied by the number of voxels in the seed mask. To correct for bias from different seed volume sizes, the waytotal was divided by the number of voxels in the seed of origin. Then, to control for directionality within the same tractograms circuit (e.g., LCL-to-ACC and ACC-to-LCL), corrected streamline counts were averaged. This averaged corrected streamline is termed ‘connectivity strength’ and has arbitrary units (a.u.). Composite group tractograms for each circuit were created by averaging each direction of the circuit for every individual, then adding all the participant images together.

### Dynamic Causal Modeling

Dynamic causal modeling (DCM; Friston et al., 2003) is a widely used technique in neuroimaging to compare different hypothetical model structures and estimate parameters describing the influence (i.e., excitatory, inhibitory, none) of one ROI on another (Stephan et al., 2010). Bayesian model comparison assigns a likelihood to each model considered.

Estimated parameters are expressed in Hz and are accompanied by posterior probabilities signifying the parameter’s strength of evidence (0.0 – 1.0: > 0.5 = “weak”, > 0.75 = “positive”, > 0.95 = “strong”, > 0.99 = “very strong”). When extracting parameters from model comparisons, a Bayesian model average weights each model by its probability, meaning average parameter estimates are influenced more by more likely models.

DCM was performed in SPM12. Prior analyses in this study modeled working memory and autobiographical memory differently. Specifically, working memory scans were modeled with 2 regressors (“working memory” and “control” conditions) while autobiographical memory scans were modeled with 4 regressors (“autobiographical onset”, “autobiographical block”, “control onset”, and “control block”). Therefore, to facilitate comparisons across tasks in the DCM, both tasks were remodeled with 2 regressors representing “experimental” trials (“working memory” from working memory and “autobiographical” from autobiographical memory) or “control” trials. These “experimental-control” models were used to derive whole-brain experimental vs. control contrast maps used in identifying circuit analysis ROIs, and to calculate experimental vs. control contrast values for each ROI in Fig. 3B.

ROI time series for ACC, PMC, SMG, and PCC were extracted from smoothed data, while LCL time series were extracted from unsmoothed, SRCC-corrected data. DCMs modeled both “experimental” and “control” conditions. Because images were not slice-time corrected, the model specification slice-timing parameter was set to the middle slice by specifying a time equal to the TR/2. DCMs used bilinear neural models, with one state per region, no stochastic effects, fit with time series, and without mean-centering of experimental input. This means experimental modulation parameters represent the change in effective connectivity in a projection from its value during the model’s implicit baseline, which here captured the ITI.

Because dMRI confirmed the presence of anatomical connections between LCL and all circuit analysis ROIs, all first level models included bidirectional inherent connectivity between LCL and ACC, PMC, SMG, and PCC, as well as inhibitory self-connections in all regions, which is a DCM default. “Experimental” condition-induced modulation of effective connectivity parameters and “experimental” condition driving input were modeled to test the hypothesis that the claustrum receives a network initiation signal and broadcasts it to task-associated regions. Specifically, the 4 circuit analysis ROIs yielded 16 models of claustrum output, comprising all logical combinations from experimental-induced modulation of claustrum projections to all 4 ROIs down to the null model, in which no claustrum projections were altered by task conditions. These 16 models were crossed with 5 possible sources of experimental input to the system: ACC, PMC, SMG, PCC, and LCL, with experimental input to LCL representing any other origin of task-related information such as primary sensory or other association cortices. This yielded 80 total hypothetical models.

A “full model” where all claustrum projections were modulated by experimental stimuli and all 5 ROIs received experimental driving input was estimated for all PIOP1 and PIOP2 working memory scans as well as all autobiographical memory scans (Zeidman et al., 2019a). All estimated first level models were compiled into a fully connected second level Parametric Empirical Bayes model (Zeidman et al., 2019b). A Bayesian model comparison was performed using the parameters of this model to assess the likelihood of all 80 hypothetical models. Bayesian model average parameters were also calculated, characterizing the task-induced change in effective connectivity of each claustrum projection by averaging the parameters across all models while weighting model parameters based on the calculated model probability. This means models deemed more likely will have a larger influence on Bayesian model average parameters than models deemed less likely.

Because comparisons of claustrum effective connectivity between memory tasks were desired, the second level design matrices contained two columns coded to specify parameters for 1) parameters describing effective connectivity in task A (either working or autobiographical memory, depending on analysis) and 2) the difference in effective connectivity between task B (the other memory task, depending on analysis) and task A. DCM task comparison results are depicted in Figures S6 (LaINS), S7 (LPulv), and S9 (LCL).

LaINS and LPulv DCMs used the same procedure except for substituting smoothed LaINS or LPulv time series for LCL time series.

### Statistical Analysis

Statistical details of experiments and the meaning of error bars are specified in figure legends. Whole-brain contrast maps were generated in SPM12. Results were voxel-wise thresholded at *p* < 0.001, followed by cluster correction to yield clusters significant at *p*-FWE < 0.05.

dMRI analyses were performed in SPSS 29.0.2.0. The Kolmogorov-Smirnov test, which is robust for samples > 50 participants (Mishra et al., 2019), was used to determine whether connectivity strengths were normally distributed. All connectivity strengths of interest had a non-normal distribution (*p* < 0.05). To compare connectivity strength with each target ROI (ACC, PMC, SMG, PCC, and Hipp) within each seed region (LCL, LaINS, and LPulv), three Related-Samples Friedman’s Two-Way ANOVA by Ranks were performed with significance set at *p* < 0.05. Dunn’s test post hoc analyses were run to determine which circuits were significantly different, with significance adjusted using Bonferroni correction for multiple comparisons. Then, five Related-Samples Friedman’s Two-Way ANOVA by Ranks were run to directly compare connectivity to the five target ROIs across seed regions with significance set to *p* < 0.05. For example, one analysis compared the LCL-ACC, LaINS-ACC, and LPulv-ACC connections. Again, Dunn’s test post hoc analyses were run with significance corrected for multiple comparisons using a Bonferroni correction. After initial Bonferroni correction, significance was set at *p* < 0.01 to account for comparisons between five ANOVAs. All *p*-values from structural connectivity analyses are recorded in Supplemental Tables 5 and 6.

DCM analyses were performed in SPM12 using SPM conventions to classify results based on posterior probability (> 0.5 = “weak”, > 0.75 = “positive”, > 0.95 = “strong”, > 0.99 = “very strong”).

All other statistical analyses were performed in GraphPad Prism 9.4.0 with significance defined as *p* < 0.05. Sidak’s multiple comparisons test was used to adjust *p*-values in ANOVA post hoc comparisons in GraphPad Prism. When performing multiple statistical tests of other kinds (e.g., multiple Spearman *r* tests), multiple comparisons correction was performed using Benjamini-Hochberg FDR correction in MATLAB.

## Figures

Structural connectivity strength comparison figures were generated in GraphPad Prism v10.3.1. All other figures were generated in GraphPad Prism v9.4.0 or MRICroGL. Group tractogram figures for LCL connections with network representative ROIs were thresholded at 50%, meaning fibers detected in at least 50% of participants are shown.

## Acknowledgments

We thank the authors of Fuentes-Claramonte et al. (2019) and Snoek et al. (2021) for sharing their data publicly on openneuro.org.

This work was supported by National Institute of Neurological Disorders and Stroke grant R01 NS112356 and National Center for Complementary and Integrative Health grant R01 AT007176, both to D.A.S.

## Author contributions

Conceptualization, B.W.S., B.N.M., and D.A.S.; Methodology, B.W.S., M.A.C., M.M., B.N.M., and D.A.S.; Formal Analysis, B.W.S., M.L.K., and M.A.C.; Investigation, B.W.S., M.L.K., and M.A.C.; Resources, M.L.K. and M.A.C.; Data Curation, M.L.K. and M.A.C.; Writing — Original Draft, B.W.S., M.A.C., B.N.M., and D.A.S.; Writing — Review & Editing, B.W.S., M.L.K., M.A.C., M.M., B.N.M., and D.A.S.; Visualization, B.W.S., M.A.C., B.N.M., and D.A.S.; Supervision: M.M., B.N.M., and D.A.S.; Project Administration, D.A.S.; Funding Acquisition, D.A.S.

## Abbreviations

ACC – Anterior Cingulate Cortex, AMI – Autobiographical Memory Interview, ANOVA – Analysis of Variance, AOMIC – Amsterdam Open MRI Collection, Autobio – Autobiographical Memory, BOLD – Blood Oxygenation Level Dependent, CSF – Cerebrospinal Fluid, DAN – Dorsal Attention Network, DCM – Dynamic Causal Modeling, DMN – Default Mode Network, dMRI – Diffusion MRI, EPI – Echo-Planar Imaging, FD – Framewise Displacement, FDR – False Discovery Rate, FOV – Field of View, FPN – Fronto-Parietal Network, FWE – Family-Wise Error, FWHM – Full Width at Half Maximum, GLM – General Linear Model, Hipp –Hippocampus, HRF – Hemodynamic Response Function, ICA – Independent Component Analysis, ITI – Inter-Trial Interval, LaINS – Left Anterior Insula, LCL – Left Claustrum, LPulv – Left Pulvinar, MNI - Montreal Neurological Institute, PCC – Posterior Cingulate Cortex, PMC – Premotor Cortex, RCL – Right Claustrum, ROI – Region of Interest, RSN – Resting State Network, SMG – Anterior Supramarginal Gyrus, SN – Salience Network, SRCC – Small Region Confound Correction, TE – Echo Time, TR – Repetition Time, VIF – Variance Inflation Factor, WM – Working Memory

