## Supplemental Material for "The human claustrum initiates networks for externally and internally driven task demands"

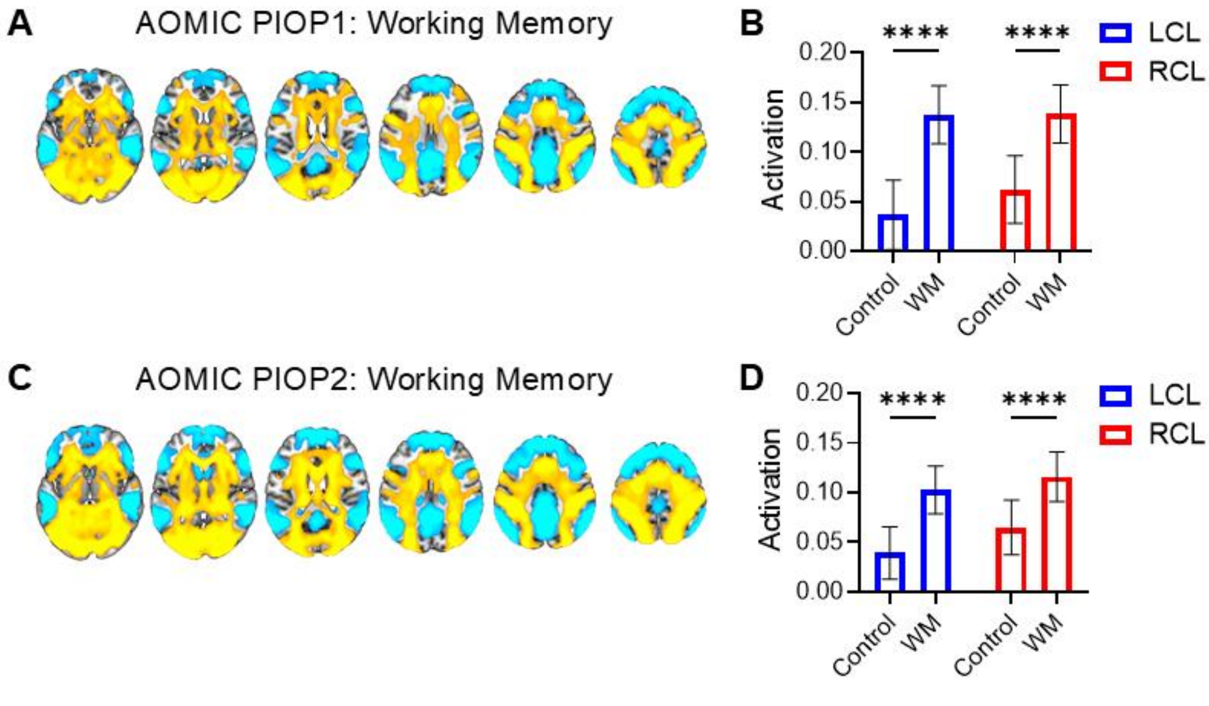


**Figure S1. Dataset-specific whole-brain and claustrum responses during working memory**

**(A)** BOLD signal increases (warm) and decreases (cool) during working memory in PIOP1 ($n$ = 198). **(B)** Average parameter estimation (regression slope) of condition-related BOLD signal change detected significantly increased bilateral claustrum activation in working memory compared to control trials in PIOP1 (two-way ANOVA main effect of condition: $F$ (1, 394) = 90.25, $p$ < 0.0001; post hoc LCL working memory vs. control: $p$ < 0.0001; post hoc RCL working memory vs. control: $p$ < 0.0001). No main effect of hemisphere ($F$ (1, 394) = 0.4013, $p$ = 0.5268) or condition x hemisphere interaction ($F$ (1, 394) = 1.787, $p$ = 0.1820) were detected. **(C)** BOLD signal increases (warm) and decreases (cool) during working memory in PIOP2 ($n$ = 222). **(D)** Two-way ANOVA detected significantly increased bilateral claustrum activation in working memory compared to control trials in PIOP2 (main effect of condition: $F$ (1, 442) = 58.74, $p$ < 0.0001; post hoc LCL working memory vs. control: $p$ < 0.0001; post hoc RCL working memory vs. control: $p$ < 0.0001). No main effect of hemisphere ($F$ (1, 442) = 1.312, $p$ = 0.2526) or condition x hemisphere interaction ($F$ (1, 442) = 0.7121, $p$ = 0.3992) were detected. Horizontal slice montages display z = 0, 10, 20, 30, 40, 50.

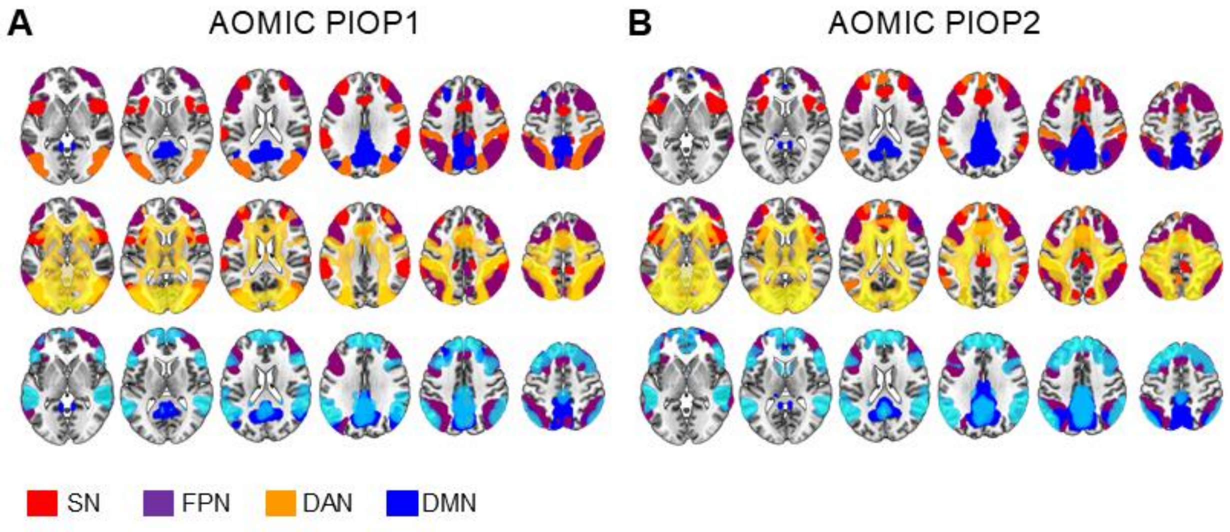


**Figure S2. Dataset-specific working memory task recruitment of resting-state networks**

**(A)** Salience (red), fronto-parietal (purple), dorsal attention (orange), and default mode (blue) networks from PIOP1 resting state group-ICA (top) with PIOP1 working memory BOLD increases (warm, middle) and decreases (cool, bottom) overlaid. **(B)** Same for PIOP2. Horizontal slice montages display z = 0, 10, 20, 30, 40, 50.

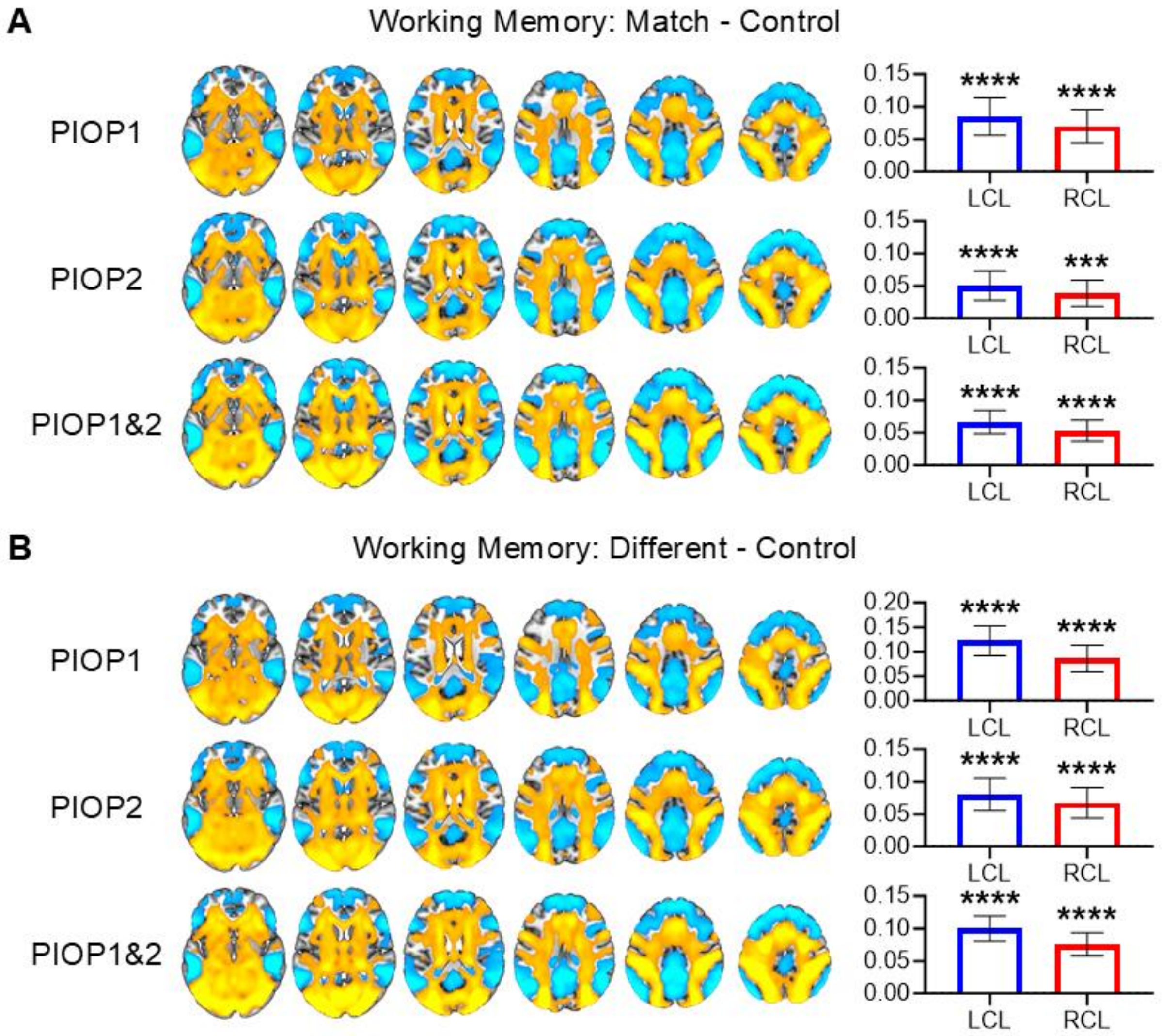


**Figure S3. “Match” and “different” working memory trials elicit bilateral network and claustrum responses.**

**(A)** BOLD signal increases (warm) and decreases (cool) from the “match - control” contrast in PIOP1, PIOP2, and the combined sample with accompanying LCL and RCL “match - control” activation (PIOP1: LCL $t = 5.745$, $p$-FDR < 0.0001; RCL $t = 5.312$, $p$-FDR < 0.0001; PIOP2: LCL $t = 4.472$, $p$-FDR < 0.0001; RCL $t = 3.754$, $p$-FDR = 0.0002; Combined: LCL $t = 7.251$, $p$-FDR < 0.0001; RCL $t = 6.447$, $p$-FDR < 0.0001). **(B)** BOLD signal increases (warm) and decreases (cool) from the “different - control” contrast in PIOP1, PIOP2, and the combined sample with accompanying LCL and RCL “different - control” activation (PIOP1: LCL $t = 7.898$, $p$-FDR < 0.0001; RCL $t = 6.172$, $p$-FDR < 0.0001; PIOP2: LCL $t = 6.439$, $p$-FDR < 0.0001; RCL $t = 5.653$, $p$-FDR < 0.0001; Combined: LCL $t = 10.13$, $p$-FDR < 0.0001; RCL $t = 8.367$, $p$-FDR < 0.0001). Horizontal slice montages display z = 0, 10, 20, 30, 40, 50.

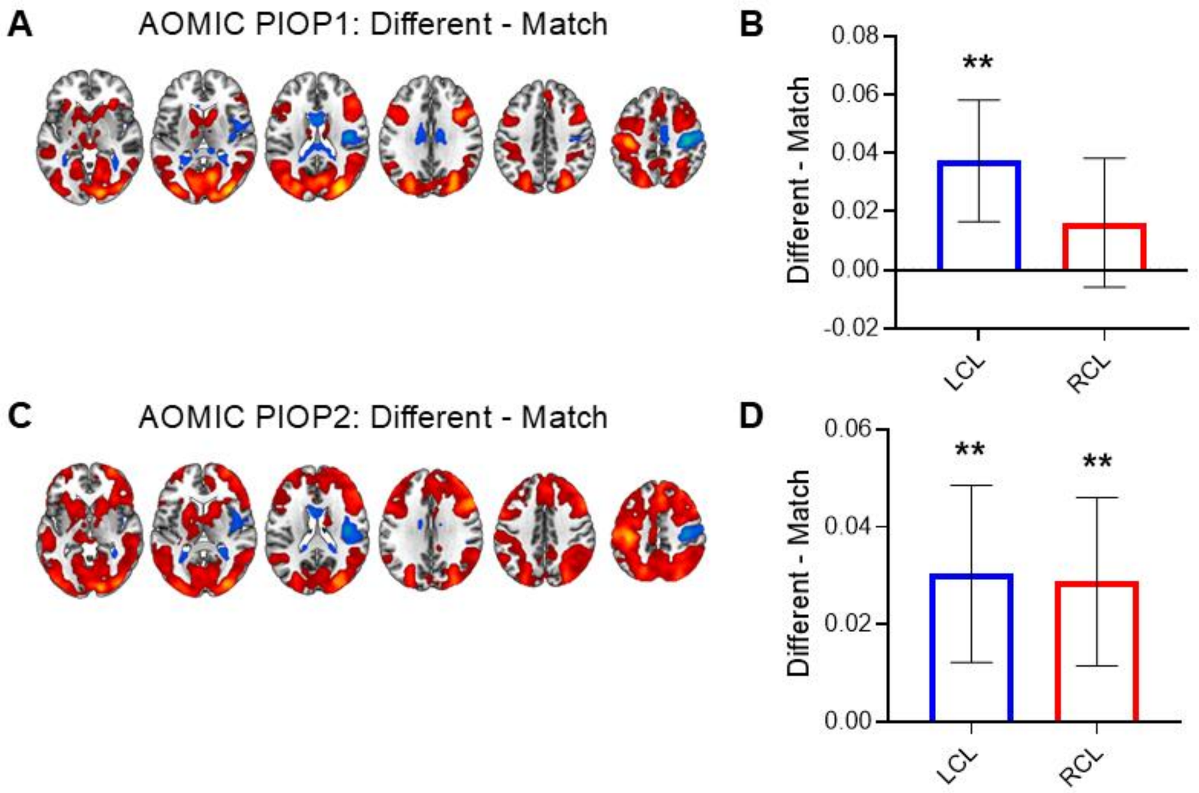


**Figure S4. Dataset-specific whole-brain and claustrum responses during different vs. match working memory trials**

**(A)** BOLD signal increases (red) and decreases (blue) from the "different - match" contrast in PIOP1. **(B)** LCL and RCL "different - match" activation in PIOP1 (LCL $t$ = 3.532, $p$-FDR = 0.0016; RCL $t$ = 1.447, $p$-FDR = 0.1496). **(C)** BOLD signal increases (red) and decreases (blue) from the "different - match" contrast in PIOP2. **(D)** LCL and RCL "different - match" activation in PIOP2 (LCL $t$ = 3.287, $p$-FDR = 0.0016; RCL $t$ = 3.274, $p$-FDR = 0.0016).

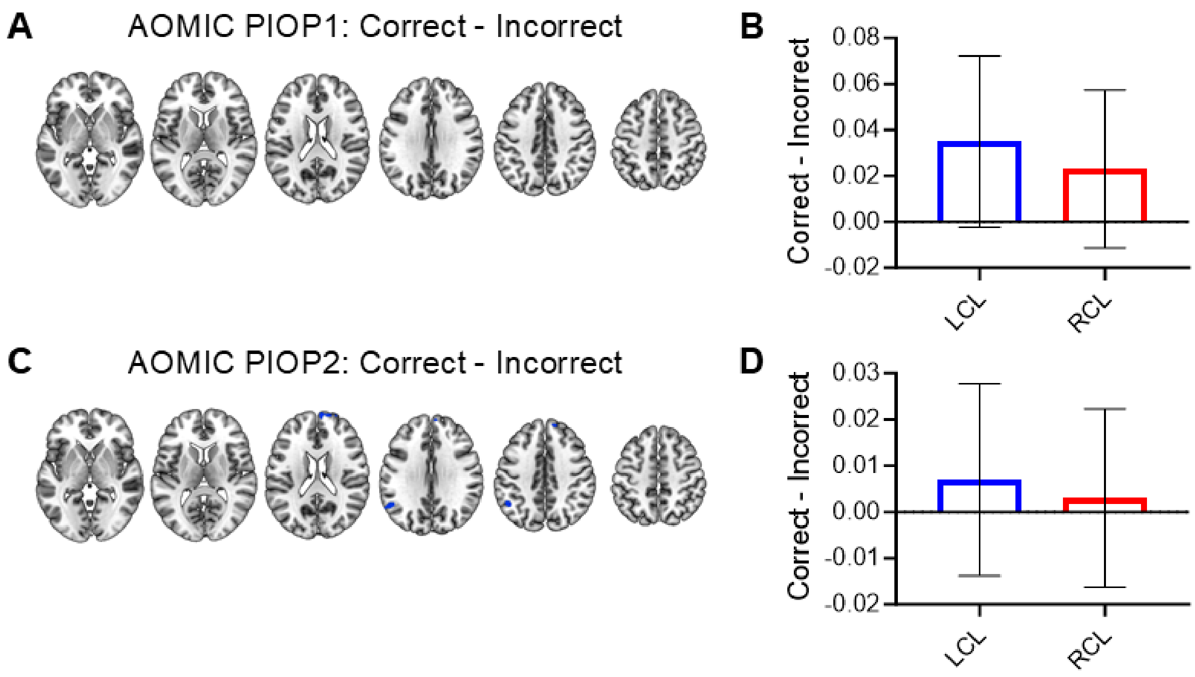


**Figure S5. Dataset-specific whole-brain and claustrum responses during correct vs. incorrect working memory trials**

**(A)** No BOLD signal increases or decreases were observed in the “correct - incorrect” contrast in PIOP1. **(B)** LCL and RCL “correct - incorrect” activation in PIOP1 (LCL $W$ = 2906, $p$ = 0.0656; RCL $W$ = 2138, $p$ = 0.1761). **(C)** No BOLD signal increases or decreases were observed in the “correct - incorrect” contrast in PIOP2. **(B)** LCL and RCL “correct - incorrect” activation in PIOP2 (LCL $W$ = 1287, $p$ = 0.4992; RCL $W$ = 711, $p$ = 0.7091).

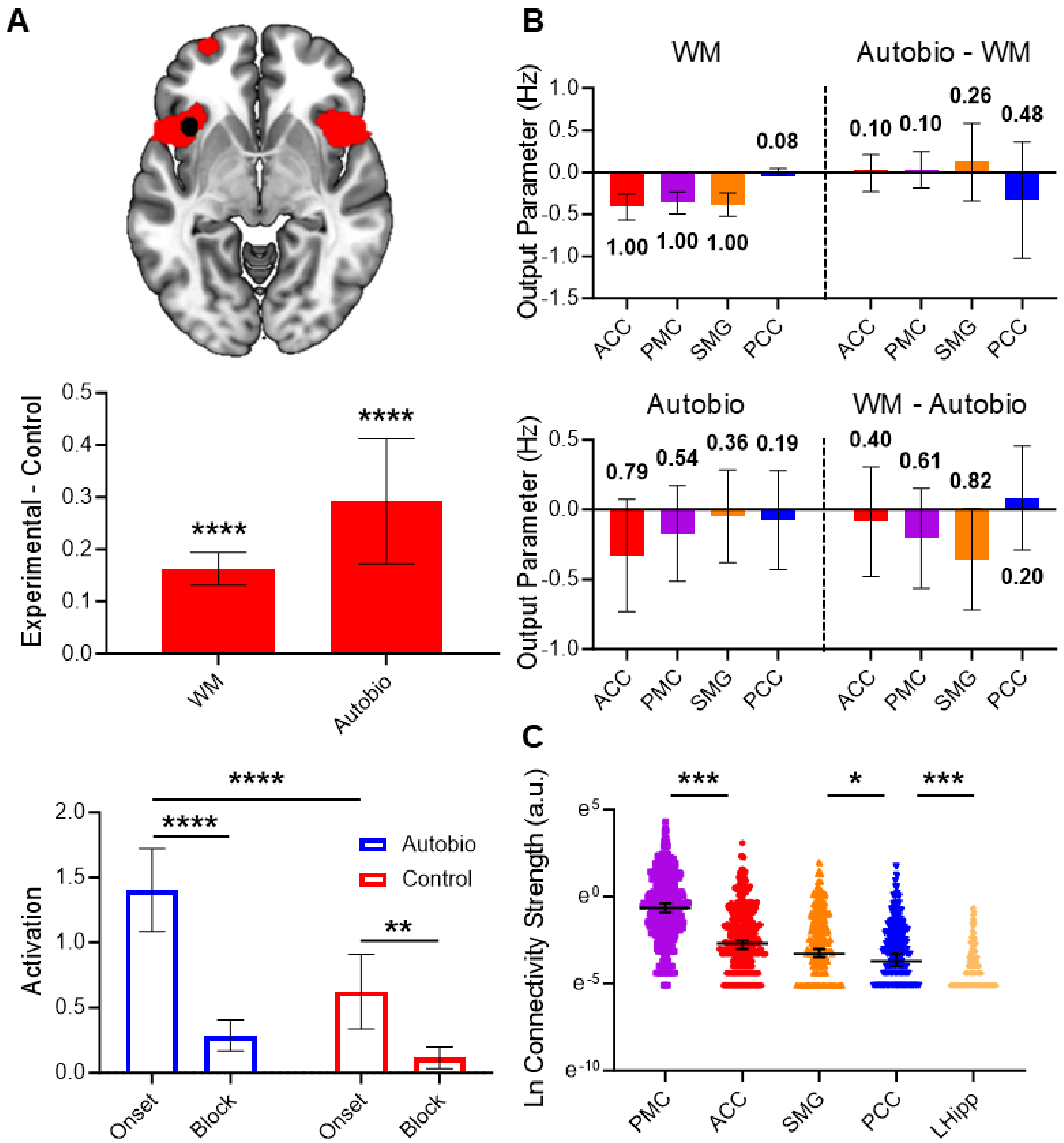


**Figure S6. Left anterior insula task response, effective connectivity, and structural connectivity profiles**

**(A) Top:** LaINS in SN (z = -5). **Middle:** "Experimental - control" contrasts revealed significant LaINS activation in WM and Autobio (WM: $t$ = 10.17, $p$-FDR < 0.0001; Autobio: $t$ = 4.95; $p$-FDR < 0.0001). **Bottom:** Hybrid event-block modeling of Autobio detected significantly increased LaINS activation during autobiographical memory and control trials (two-way ANOVA main effect of condition: $F$ (1, 68) = 18.08, $p$ < 0.0001; main effect of timepoint: $F$ (1, 68) = 54.64, $p$ < 0.0001; condition x timepoint interaction: $F$ (1, 68) = 7.599, $p$ = 0.0075; post hoc Autobio onset vs. Autobio block: $p$ < 0.0001; post hoc control onset vs. control block: $p$ = 0.0033; post hoc Autobio onset vs. control onset: $p$ < 0.0001). **(B) Top:** Estimated output parameters during WM and differences with Autobio. **Bottom:** Estimated output parameters during Autobio and differences with WM. **(C)** LaINs exhibited preferential structural connectivity with PMC. Select comparisons shown for clarity.

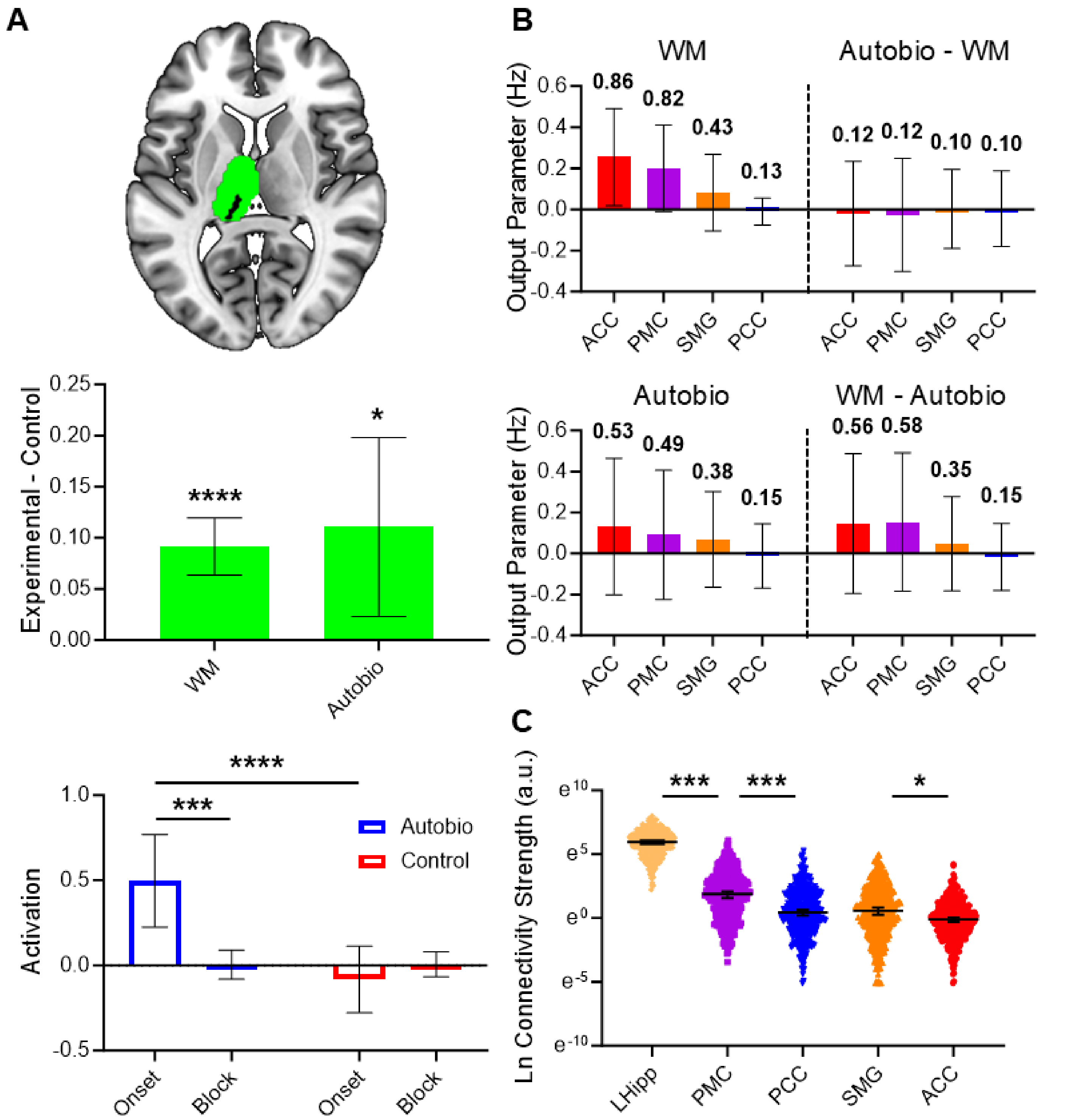


**Figure S7. Left pulvinar task response, effective connectivity, and structural connectivity profiles**

**(A) Top:** LPulv in left thalamus (Cole et al., 2019; z = 8). **Middle:** "Experimental - control" contrasts revealed significant LPulv activation in WM and Autobio tasks (WM: $t = 6.47$, $p$-FDR $< 0.0001$; Autobio: $t = 2.57$; $p$-FDR $= 0.0149$). **Bottom:** Hybrid event-block modeling of Autobio detected significantly increased LPulv activation at autobiographical memory trial onset (two-way ANOVA main effect of condition: $F$ (1, 68) = 13.25, $p < 0.0005$; main effect of timepoint: $F$ (1, 68) = 4.640, $p < 0.0348$; condition x timepoint interaction: $F$ (1, 68) = 9.595, $p = 0.0028$; post hoc Autobio onset vs. Autobio block: $p$ = 0.0008; post hoc Autobio onset vs. control onset: $p < 0.0001$). However, Autobio onset activation was positively correlated with subject motion (Supplemental Table 9). **(B) Top:** Estimated output parameters during WM and differences with Autobio. **Bottom:** Estimated output parameters during Autobio and differences with WM. **(C)** LPulv exhibited preferential structural connectivity with Hipp. Select comparisons shown for clarity.

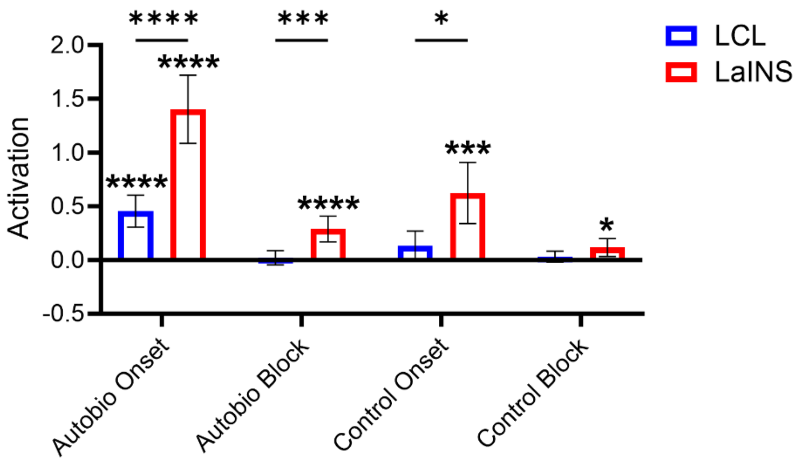


**Figure S8. Larger, longer, and less specific LaINS than LCL responses to autobiographical memory scan stimuli**

One sample *t*-tests of LaINS and LCL BOLD responses to autobiographical memory scan stimuli detected significant activation above baseline in LaINS to all conditions, but only to Autobio onset in LCL (LaINS: Autobio onset $t = 8.974$, *p*-FDR < 0.0001; Autobio block $t = 4.907$, *p*-FDR < 0.0001; Control onset $t = 4.455$, *p*-FDR = 0.0002; Control block $t = 2.806$, *p*-FDR = 0.0132; LCL: Autobio onset $t = 6.194$, *p*-FDR < 0.0001; Autobio block $t = 0.6129$, *p*-FDR = 0.5440; Control onset $t = 1.936$, *p*-FDR = 0.0817; Control block $t = 1.249$, *p*-FDR = 0.2516). Two-way ANOVA detected significantly greater LaINS than LCL signal (main effect of ROI: $F(1, 68) = 40.71$, $p < 0.0001$; main effect of condition: $F(2.086, 141.9) = 43.46$, $p < 0.0001$; ROI x condition interaction: $F(3, 204) = 10.20$, $p < 0.0001$), with post hoc comparisons identifying greater LaINS than LCL signal during Autobio onset ($p < 0.0001$), Autobio block ($p = 0.0008$), and Control onset ($p = 0.0104$), but not Control block ($p = 0.3049$).

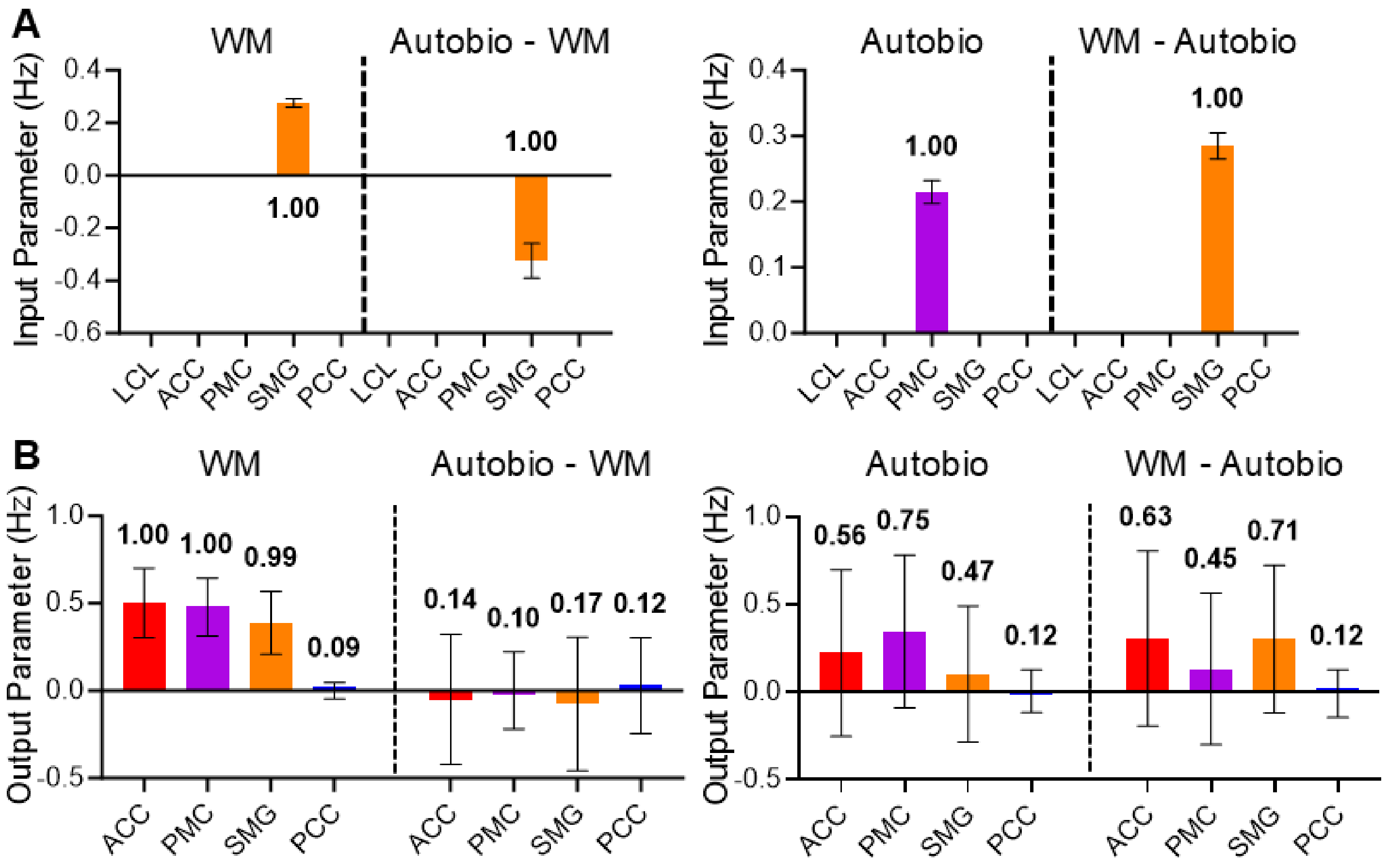


**Figure S9. Left claustrum DCM task comparisons**

**(A) Left:** Estimated input parameters during WM and differences with Autobio. **Right:** Estimated input parameters during Autobio and differences with WM. **(B) Left:** Estimated output parameters during WM and differences with Autobio. **Right:** Estimated output parameters during Autobio and differences with WM.

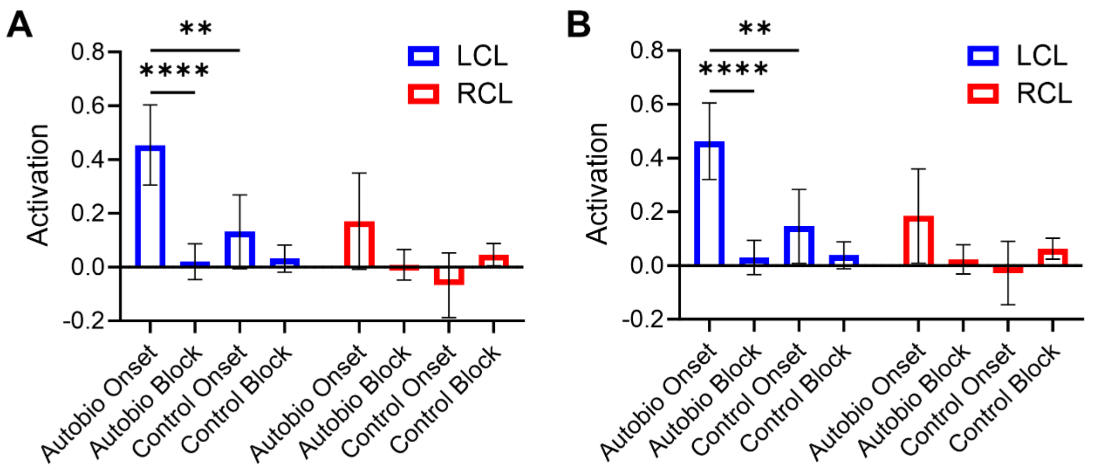


**Figure S10. Claustrum results were not materially impacted by removal of the first 10 volumes.**

**(A)** LCL and RCL activation during autobiographical memory scans, reproduced from Fig. 2C. **(B)** LCL and RCL activation during autobiographical memory scans following preprocessing which included removal of the initial 10 volumes. Bilateral claustrum results are qualitatively similar (two-way ANOVA main effect of condition: $F$ (2.149, 146.2) = 13.69, $p < 0.0001$; main effect of hemisphere: $F$ (1, 68) = 8.080, $p = 0.0059$; condition x hemisphere interaction: $F$ (3, 204) = 3.559, $p = 0.0152$; post hoc LCL Autobio onset vs. Autobio block: $p < 0.0001$; post hoc LCL Autobio onset vs. control onset: $p = 0.0056$).

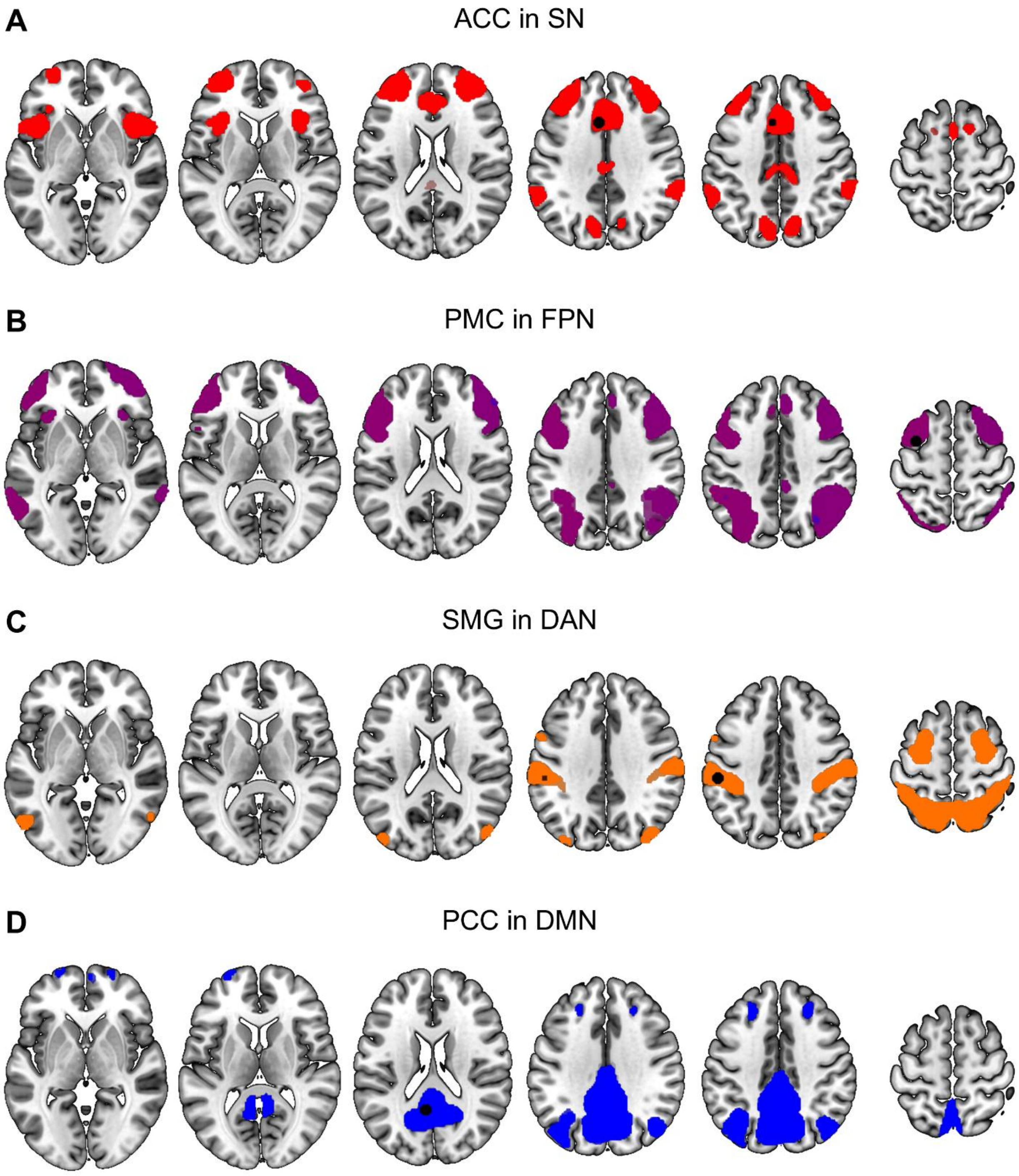


**Figure S11. Network representative node ROIs overlaid on isolated resting-state networks from combined PIOP1 & PIOP2 group-ICA**

Network representative node ROIs (black circles) overlaid on RSNs for better visualization. **(A)** ACC in SN. **(B)** PMC in FPN. **(C)** SMG in DAN. **(D)** PCC in DMN.

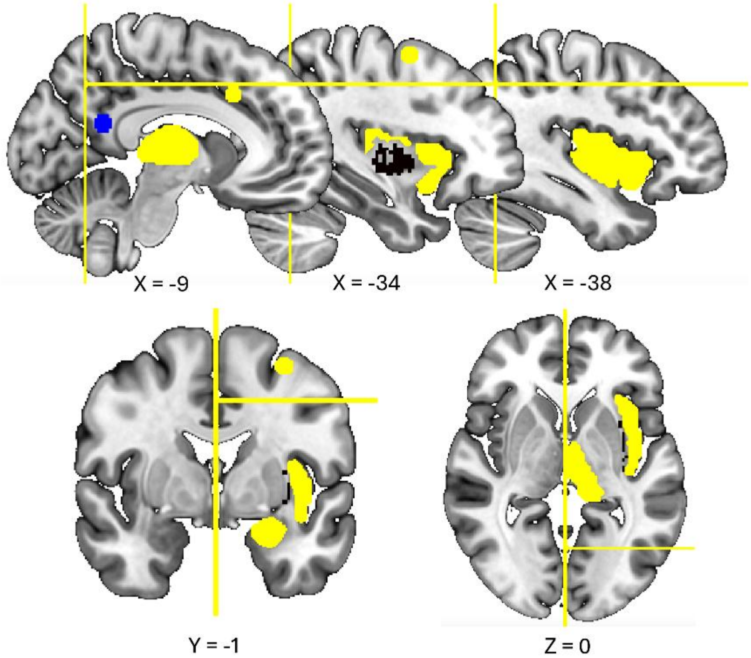


**Figure S12. Representative dMRI analysis exclusion masks**

Representative structural connectivity analysis exclusion mask (yellow) setup for the connection between LCL (black, top middle) and PCC (blue, top left). Analyses used combinations of unanalyzed target ROIs, nearby notable ROIs, and additional planes for specificity. LCL-PCC analyses used ROI exclusion masks for ACC (top left), PMC (top middle, bottom left), SMG, and Hipp, as well as masks for thalamus, insular cortex, and amygdala. Exclusion planes for LCL-PCC included a mid-sagittal plane (x = 0), a horizontal plane at z = 42, and a coronal plane at y = -68. Images displayed in radiological orientation.

| Task & Dataset | LCL-AVG FD | RCL-AVG FD |
|---|---|---|
| Working Memory (PIOP1)<br>WM vs. baseline | *r* = -0.01568<br>*p* = 0.8265<br>*p*-FDR = 0.8449 | *r* = -0.1578<br>*p* = 0.0264*<br>*p*-FDR = 0.1584 |
| Working Memory (PIOP2)<br>WM vs. baseline | *r* = -0.01321<br>*p* = 0.8449<br>*p*-FDR = 0.8449 | *r* = 0.02156<br>*p* = 0.7493<br>*p*-FDR = 0.8449 |
| Autobiographical Memory<br>Autobio onset vs. baseline | *r* = 0.05602<br>*p* = 0.7492<br>*p*-FDR = 0.8449 | *r* = 0.2793<br>*p* = 0.1042<br>*p*-FDR = 0.3126 |

**Supplemental Table 1. No significant positive correlations between subject motion and claustrum BOLD signal change.**

Spearman *r* tests (average framewise displacement [FD] values were not normally distributed) find no significant correlations between individual participant LCL or RCL BOLD signal activation parameters and average motion during task scans. Working memory correlations used the working memory condition vs. implicit baseline contrast, and autobiographical memory correlations used the autobiographical memory onset condition vs. implicit baseline contrast. Implicit baseline in both models encompassed inter-trial intervals (ITIs) and volumes obtained prior to and following task runs. Only RCL-AVG FD during PIOP1 working memory yielded an uncorrected *p*-value less than 0.05 ($r = -0.1578$; $p < 0.0264$), but the inverse correlation revealed RCL signal decreased with greater motion, meaning motion-induced noise potentially biased observations away from significant condition effects.

| | All Working Memory | | | All Control | | Working Memory: Different | | | Working Memory: Match | | |
|---|---|---|---|---|---|---|---|---|---|---|---|
| Dataset | Correct | Incorrect | Miss | Hit | Miss | Correct | Incorrect | Miss | Correct | Incorrect | Miss |
| PIOP1 | 53% | 21% | 26% | 49% | 51% | 55% | 18% | 28% | 51% | 25% | 24% |
| PIOP2 | 51% | 47% | 2% | 73% | 27% | 52% | 46% | 2% | 50% | 48% | 2% |

**Supplemental Table 2. Working memory task accuracy by trial type.**

Because accuracy percentages were not normally distributed, Wilcoxon matched-pairs signed rank tests were performed to compare accuracies between conditions. “Different” and “match” accuracies were not significantly different in either dataset (PIOP1: $p = 0.0797$; PIOP2: $p = 0.1815$).

| | All Working Memory | | All Control | Working Memory: Different | | Working Memory: Match | |
|---|---|---|---|---|---|---|---|
| Dataset | Correct | Incorrect | Hit | Correct | Incorrect | Correct | Incorrect |
| PIOP1 | 0.57s | 0.58s | 0.68s | 0.58s | 0.56s | 0.57s | 0.59s |
| PIOP2 | 0.45s | 0.47s | 0.88s | 0.45s | 0.48s | 0.44s | 0.46s |

**Supplemental Table 3. Working memory task mean reaction time by trial type.**

Mixed Effects Analysis did not detect a significant effect of condition (different vs. match) or accuracy (correct vs. incorrect) on mean reaction time in either PIOP1 or PIOP2. Only a condition x accuracy interaction was detected in PIOP1 ($p$ = 0.0132), where incorrect responses were made faster than correct responses in “different” trials.

| Task & Dataset | LaINS-AVG FD | LPulv-AVG FD |
| --- | --- | --- |
| Working Memory (PIOP1)<br>WM vs. baseline | *r* = -0.01522<br>*p* = 0.8314<br>*p*-FDR = 0.9953 | *r* = -0.1326<br>*p* = 0.0626<br>*p*-FDR = 0.1252 |
| Working Memory (PIOP2)<br>WM vs. baseline | *r* = 0.006146<br>*p* = 0.9274<br>*p*-FDR = 0.9953 | *r* = 0.0003965<br>*p* = 0.9953<br>*p*-FDR = 0.9953 |
| Autobiographical Memory<br>Autobio onset vs. baseline | *r* = 0.3258<br>*p* = 0.0562<br>*p*-FDR = 0.1252 | *r* = 0.5280<br>*p* = 0.0011**<br>*p*-FDR = 0.0066** |

**Supplemental Table 4. Correlations between subject motion and LaINS and LPulv BOLD signal change.**

Spearman *r* tests (average framewise displacement [FD] values were not normally distributed) find no significant correlations between individual participant LaINS BOLD signal activation and average motion during task scans. However, a significant positive correlation was found between LPulv BOLD signal activation and average motion during autobiographical memory scans ($r = 0.5280$, $p$-FDR = 0.0066), calling into question the validity of the observed task-induced BOLD increase in the region. Working memory correlations used the working memory condition vs. implicit baseline contrast, and autobiographical memory correlations used the autobiographical memory onset condition vs. implicit baseline contrast. Implicit baseline in both models encompassed inter-trial intervals (ITIs) and volumes obtained prior to and following task runs.

| Within Seeds | Connection | Standardized Test Statistic (*z*) | Adjusted Significance (*p*) |
|---|---|---|---|
| LCL | PCC-Hipp | 6.83 | <0.001 |
| | ACC-Hipp | 8.751 | <0.001 |
| | SMG-Hipp | 21.713 | <0.001 |
| | PMC-Hipp | 28.281 | <0.001 |
| | ACC-PCC | 1.92 | 0.548 |
| | SMG-PCC | 14.882 | <0.001 |
| | PMC-PCC | 21.451 | <0.001 |
| | SMG-ACC | 12.962 | <0.001 |
| | PMC-ACC | 19.531 | <0.001 |
| | PMC-SMG | 6.568 | <0.001 |
| LaINS | PCC-Hipp | 9.438 | <0.001 |
| | SMG-Hipp | 12.482 | <0.001 |
| | ACC-Hipp | 14.948 | <0.001 |
| | PMC-Hipp | 25.488 | <0.001 |
| | SMG-PCC | 3.044 | 0.023 |
| | ACC-PCC | 5.51 | <0.001 |
| | PMC-PCC | 16.05 | <0.001 |
| | ACC-SMG | 2.466 | 0.137 |
| | PMC-SMG | 13.006 | <0.001 |
| | PMC-ACC | 10.54 | <0.001 |
| LPulv | SMG-ACC | 3.11 | 0.019 |
| | PCC-ACC | 4.015 | 0.001 |
| | PMC-ACC | 11.009 | <0.001 |
| | Hipp-ACC | 26.874 | <0.001 |
| | PCC-SMG | 0.906 | 1.000 |
| | PMC-SMG | 7.899 | <0.001 |
| | Hipp-SMG | 23.764 | <0.001 |
| | PMC-PCC | 6.994 | <0.001 |
| | Hipp-PCC | 22.858 | <0.001 |
| | Hipp-PMC | 15.864 | <0.001 |

**Supplemental Table 5. Within seed ROI structural connectivity post hoc comparison *p*-values.**

All *p*-values are adjusted using Bonferroni correction within each seed analysis.

| Target | Between Seeds | Standardized Test Statistic (*z*) | Adjusted Significance (*p*) |
|---|---|---|---|
| ACC | LPulv-LaINS | 15.768 | <0.001 |
| | LCL-LaINS | 20.978 | <0.001 |
| | LCL-LPulv | 5.21 | <0.001 |
| PMC | LPulv-LaINS | 10.144 | <0.001 |
| | LCL-LaINS | 24.118 | <0.001 |
| | LCL-LPulv | 13.974 | <0.001 |
| SMG | LPulv-LaINS | 13.422 | <0.001 |
| | LCL-LaINS | 26.43 | <0.001 |
| | LCL-LPulv | 13.008 | <0.001 |
| PCC | LCL-LaINS | 18.25 | <0.001 |
| | LPulv-LaINS | 20.202 | <0.001 |
| | LPulv-LCL | 1.949 | 0.154 |
| Hipp | LCL-LaINS | 14.04 | <0.001 |
| | LPulv-LaINS | 28.76 | <0.001 |
| | LPulv-LCL | 14.716 | <0.001 |

**Supplemental Table 6. Between seed ROI structural connectivity post hoc comparison *p*-values.**

All *p*-values are adjusted using Bonferroni correction within each seed analysis. Because comparisons include 5 separate ANOVAs, significance is set at $p < 0.05/5$, or $p < 0.01$.

| ROI | Task | 1 | 2 | 3 | 4 | 5 | 6 | 7 | 8 | 9 | 10 | 11 | 12 | 13 | 14 | 15 | 16 |
|---|---|---|---|---|---|---|---|---|---|---|---|---|---|---|---|---|---|
| LCL | WM | 0.09 | 0.90 | 0.00 | 0.01 | 0.00 | 0.00 | 0.00 | 0.00 | 0.00 | 0.00 | 0.00 | 0.00 | 0.00 | 0.00 | 0.00 | 0.00 |
| | Autobio - WM | 0.00 | 0.00 | 0.00 | 0.01 | 0.00 | 0.02 | 0.01 | 0.09 | 0.00 | 0.01 | 0.01 | 0.07 | 0.02 | 0.11 | 0.08 | 0.56 |
| | Autobio | 0.02 | 0.18 | 0.03 | 0.20 | 0.01 | 0.06 | 0.01 | 0.07 | 0.02 | 0.14 | 0.02 | 0.15 | 0.01 | 0.05 | 0.01 | 0.05 |
| | WM - Autobio | 0.02 | 0.18 | 0.01 | 0.07 | 0.03 | 0.22 | 0.01 | 0.09 | 0.01 | 0.10 | 0.01 | 0.04 | 0.02 | 0.13 | 0.01 | 0.05 |
| LaINS | WM | 0.08 | 0.92 | 0.00 | 0.00 | 0.00 | 0.00 | 0.00 | 0.00 | 0.00 | 0.00 | 0.00 | 0.00 | 0.00 | 0.00 | 0.00 | 0.00 |
| | Autobio - WM | 0.00 | 0.00 | 0.00 | 0.00 | 0.01 | 0.01 | 0.03 | 0.03 | 0.01 | 0.01 | 0.03 | 0.04 | 0.10 | 0.11 | 0.29 | 0.32 |
| | Autobio | 0.03 | 0.12 | 0.05 | 0.22 | 0.03 | 0.11 | 0.04 | 0.19 | 0.01 | 0.03 | 0.01 | 0.06 | 0.01 | 0.03 | 0.01 | 0.05 |
| | WM - Autobio | 0.04 | 0.16 | 0.01 | 0.03 | 0.03 | 0.10 | 0.01 | 0.02 | 0.06 | 0.24 | 0.01 | 0.05 | 0.04 | 0.15 | 0.01 | 0.03 |
| LPulv | WM | 0.04 | 0.26 | 0.05 | 0.34 | 0.01 | 0.06 | 0.01 | 0.08 | 0.01 | 0.04 | 0.01 | 0.06 | 0.00 | 0.01 | 0.00 | 0.02 |
| | Autobio - WM | 0.00 | 0.00 | 0.00 | 0.01 | 0.00 | 0.01 | 0.01 | 0.08 | 0.00 | 0.01 | 0.01 | 0.08 | 0.01 | 0.07 | 0.07 | 0.63 |
| | Autobio | 0.01 | 0.08 | 0.02 | 0.14 | 0.02 | 0.09 | 0.03 | 0.14 | 0.01 | 0.07 | 0.02 | 0.12 | 0.01 | 0.08 | 0.02 | 0.13 |
| | WM - Autobio | 0.02 | 0.10 | 0.03 | 0.18 | 0.01 | 0.07 | 0.02 | 0.13 | 0.01 | 0.08 | 0.03 | 0.14 | 0.01 | 0.05 | 0.02 | 0.10 |

**Supplemental Table 7. Output Bayesian model comparison probabilities**

Probabilities assigned to models with different combinations of task-modulation of network representative node outputs. Values reveal which model structure is favored, but they do not provide insight into the parameters (e.g., excitatory or inhibitory) describing the models.

| | Active | | | | |
|---|---|---|---|---|---|
| Regressor | Alert | Encoding | Retention | Probe | Response |
| VIF | 149.18 | 640.43 | 675.06 | 443.29 | 82.17 |
| | Passive | | | | |
| Regressors | Alert | Encoding | Retention | Probe | Response |
| VIF | 257.74 | 1215.50 | 1345.83 | 824.19 | 138.07 |

**Supplemental Table 8. High likelihood of misattribution in working memory GLMs with individual task component regressors.**

Variance Inflation Factors (VIFs) reveal high likelihood of misattribution when modeling each task component (alert, encoding, retention, test, response) of active and passive working memory conditions. Common "rules of thumb" regard VIFs exceeding 5 or 10 as suggesting high levels of multicollinearity that can severely impact model stability (O'Brien, 2007).

| Regressor | Autobio Onset | Autobio Block | Control Onset | Control Block |
|---|---|---|---|---|
| VIF | 1.39 | 1.44 | 1.37 | 1.44 |

**Supplemental Table 9. No risk of misattribution in autobiographical memory GLMs when using hybrid event-block format.**

All VIFs using hybrid event-block format in autobiographical memory task GLMs are less than 2, reflecting no risk of model instability or misattribution.
